# Evolution of Species’ Range and Niche in Changing Environments

**DOI:** 10.1101/2025.01.16.633367

**Authors:** Jitka Polechová

## Abstract

What causes species’ niche and range margins to shift is not only a fundamental theoretical question, but also directly affects how we assess the resilience of natural populations in current and future environments. Yet despite the urgent need for theory that can predict evolutionary and ecological responses in times of accelerated climate change, the assumptions of current eco-evolutionary theory remain restrictive, with predictions neglecting important interactions between ecological and evolutionary forces. In this study, I provide quantitative, testable predictions on limits to adaptation in changing environments, which arise from the feedback between selection, genetic drift, and population dynamics. This eco-evolutionary feedback creates a tipping point beyond which adaptation fails as genetic drift overwhelms selection, and species’ ranges contract from the margins or fragment abruptly – even under gradual environmental change. This “expansion threshold” is determined by three parameters: two quantifying the effects of spatial and temporal variability on the fitness of the population, and one capturing the impact of genetic drift: the reduction of local genetic diversity across generations in finite populations. Genetic drift is strong in populations with small “neighbourhood size”. Increasing dispersal, such as via assisted migration, enlarges neighbourhood size, counteracting the loss of genetic variation due to genetic drift. This increases adaptive potential and can facilitate evolutionary rescue in changing environments. Conversely, beyond the expansion threshold, local genetic variance becomes depleted, increasing extinction probability. The theory provides general predictions for species’ range and niche dynamics beyond standard ecological niche models, highlighting the fundamental impact of eco-evolutionary interactions.

**Significance Statement:** In a time of accelerating climate change, we need a predictive theory of species’ range shifts, adaptation, and resilience. Currently, predictions rely on theory that fails to incorporate the effects of interactions between ecological and evolutionary forces. This study demonstrates that eco-evolutionary feedback between selection, genetic drift, and population dynamics can create a tipping point at which genetic variance becomes depleted and species’ ranges contract or fragment. This tipping point depends on three parameters: two capturing loss of fitness due to spatial and temporal change, and one quantifying genetic drift, increasing as population density and dispersal decrease. By advancing the theoretical foundations of species’ range dynamics in changing environments, this study can inform the development of strategies in conservation genetics.

## Introduction

As environments change over time, species’ ranges may shift, expand, or contract. These ecological responses are profoundly influenced by evolutionary processes: changes in mean fitness drive changes in population size, while variance in fitness drives adaptation (1–4). Although theoretical progress has been made in understanding range shifts, this progress has come at a cost of significant simplification. In particular, most current theory assumes fixed genetic variance (5–8), even though genetic variance evolves under eco-evolutionary feedback, fundamentally changing the expected limits to adaptation and species’ range limits.

When environments vary through both time and space, their effects on fitness and genetic variation interact, influencing adaptation rates and the probability of evolutionary rescue (9–14). In spatially structured populations, a substantial proportion of genetic variance can be maintained by gene flow (15–17) – and spatial subdivision can substantially extend the time scales over which recurrent selective sweeps elevate local genetic variance, relative to well-mixed populations (18, 19).

To date, however, eco-evolutionary studies accounting for evolving genetic variation in jointly spatially and temporally varying environments have only rarely been attempted (see (20–23)) and we lack a framework for predictive, quantitative results on the evolution of species’ range and niche in realistic, spatially and temporally changing environments that include the effects of genetic drift.

Critical to developing such theoretical framework is accurate accounting for the effect of dispersal of individuals through space, long recognised as a an important factor for species’ range limits as well as limits to adaptation in general (24, 25). Population subdivision and dispersal play a central role in Wright’s (26) shifting balance theory, which posits that increased drift in subdivided populations can facilitate crossing fitness valleys. However, dispersal can fundamentally affect adaptation even on smooth fitness landscapes. Already Haldane (27) introduced the idea that asymmetric gene flow from the centre to the edges of a species’ range could swamp local adaptation at the margin, confining the species to a defined range, an example of migrational meltdown (28). On the other hand, dispersal can promote expansion of species’ ranges, through elevating adaptive genetic variance (29, 30), which enables adaptation (31, 32). The conflicting qualitative predictions (15, 24, 33) imply a need for explicit models which include the feedback between population size and adaptation as well as evolution of genetic variance.

Dispersal also has an important effect on demographic fluctuations and genetic stochasticity. By enlarging the neighbourhood size, which can be thought of as a local mating pool, dispersal reduces genetic drift, i.e., stochastic fluctuations in allele frequencies. Neighbourhood size was first defined by Wright (34) as the size of a single panmictic population that would give the same probability of identity-by-descent in the previous generation, and quantifies the strength of genetic drift in populations distributed across continuous space. More generally, it can be defined as the ratio of the rate at which individuals spread through space to the per-generation rate of coalescence (35). In two dimensions with diffusive dispersal, the neighbourhood size then corresponds to 𝒩= 4*π σ*^2^ *N*, a population within the area of a circle with a radius 2*σ* and local (effective) population density *N* . In a haploid population, we get 𝒩 = 2*π σ*^2^ *N* .

Neighbourhood size 𝒩 reflects the local strength of genetic drift, which is particularly relevant for diversifying selection. In two-dimensional habitats, 𝒩 is dimensionless. In contrast, effective population size *N*_*e*_ has units of time and reflects the rate of genetic drift in a sampled population. Population-wide (total) effective population size *N*_*e,T*_ may either increase or decrease with dispersal: *N*_*e,T*_ decreases with dispersal when local population density fluctuations are low or absent, but can increase if dispersal dampens large demographic fluctuations (16, 36, 37). Local effective population size *N*_*e,L*_ increases with dispersal at equilibrium; however, this effect can be reversed over short timescales when allele frequencies change faster than expected due to a bout of dispersal, mimicking genetic drift (38). Interestingly, although a sufficiently large neighbourhood size is critical for the maintenance of adaptive diversity across space (16, 39, 40), it is – unlike effective population size or census size – currently almost never explicitly considered in conservation guidelines [though see (41–43)]. Despite a growing understanding that higher dispersal and gene flow may enhance population resilience, especially under changing environments (44), explicit consideration of population structure, and neighbourhood size in particular, remains neglected.

Here, I develop a framework for predictive quantitative results of range and niche evolution in realistic, spatially and temporally changing environments. This model incorporates five dimensionless parameters capturing rates of environmental change in space and time, population size and gene flow, as well as selection and mutation, allowing predictions of species’ range collapse or expansion. These predictions highlight an important feedback between ecological and evolutionary processes, leading to a tipping point where genetic drift overwhelms adaptation.

### Model and methodology summary

The core features of the model are eco-evolutionary feedback through the coupling between fitness and population growth (Eq. 1 and 2), the inclusion of both genetic and demographic stochasticity, and a quantitative genetic model in which both the trait mean and its variance evolve via change in underlying allele frequencies (Eq. 3 and 4). The effect of environment (biotic and abiotic) is represented by a gradient in the optimum value of a trait of the focal species. The optimum changes through time along one spatial coordinate: an example could be a spatial gradient in optimal flowering time, changing with altitude (but not along the contours of a hill) and time. Both range and niche shifts are possible, as suitable habitats can be accessed through dispersal, while the selected trait – including its variance – evolves in response to environmental change.

Individual fitness depends on both the local density and the distance of the trait from the environmental optimum. The population growth is given by the mean fitness, which reflects the local density and phenotypic distance from the optimum (see Eq. 1). Selection is hard, i.e., the locally attainable density (effective carrying capacity’) declines with maladaptation. If selection were acting on relative fitness only (soft selection *sensu* Christiansen (45)) and the carrying capacity were fixed, there would be no range margins unless they were *a priori* imposed by niche vacancies to be filled. Note that Christiansen’s definition differs from how hard and soft selection were originally defined by Wallace (46), motivated by the problem of the maintenance of adaptive variation when selection carries a demographic cost (4, 47). We say that the selection is hard (*sensu* (45)) in our model, as the population growth is reduced due to maladaptation, while the regulation of local population size is density-dependent (Eq. 2). Yet, the allele frequencies underlying the polygenic trait are under frequency-dependent selection, and there is no density-dependent selection (Eq. 4).

The additive quantitative trait that determines adaptation to the changing environment is determined by a large number of loci with exponentially distributed effects *α*_*i*_. There is free recombination and for simplicity, individuals are assumed to be hermaphroditic and haploid. Thus, there are no dominance effects; yet there is epistasis as the fitness effect of each allele depends on other allele frequencies via the trait mean. The dispersal distances are normally distributed, independent of fitness and do not evolve through time.

Building on fundamental population genetics theory (1, 3, 48) as well as preceding evolutionary-genetic theory of species’ ranges (6, 49–51), the model is written both as a system of coupled stochastic partial differential equations (SPDEs) in continuous time and spacer as well as an individual-based discrete-time model (IBM) with space represented by a two-dimensional lattice and a simple yet explicit genetic architecture. The IBM, representing a more complex eco-evolutionary representation, constitutes one of many examples approximated by the SPDEs.

The eco-evolutionary dynamics is elucidated with the help of the following steps. First, the SPDE formalisation in continuous time provides a rescaling in terms of dimensionless parameters, which reduces the apparent dimensionality of the system. Second, the outcomes of the IBM are compared to known analytical expectations of simpler systems: namely to the deterministic equilibria and the predictions of a phenotypic model with fixed genetic variance. Third, the robustness of the predictive parameters arising from the non-dimensionalisation is then tested using the IBM. An illustrative example of the simulated IBM is given at Fig. S1.

## Results

### Scale-free parameters of the model

This model of species’ range and niche dynamics has over ten parameters, which arise from coupling of evolutionary and ecological dynamics (Eq. 1–4). Fortunately, there is a rescaling which both reduces the apparent dimensionality of the system and facilitates interpretation. Upon nondimensionalization, we obtain five dimensionless parameters that describe the core system – see Eq. 5 and Table 1, extending from (6, 51, 52). The first parameter measures the effective rate of change of the optimum through time: the rate of temporal change *k* is scaled as 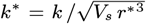, relative to the strength of stabilising selection *V*_*s*_ and the rate 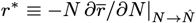 at which the population returns to its demographic equilibrium 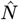 upon a small perturbation. The second parameter, 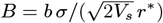, describes the effective steepness of the environmental gradient *b* across space *x* in terms of the expected reduction in fitness due to dispersal *σ*. The third parameter is the (haploid) neighbourhood size 𝒩 = 2*π σ*^2^ *N*, where *N* is the local population density. 𝒩 describes the population attainable within a generation’s dispersal and measures the strength of genetic drift on local scales, scaling with 1*/*𝒩 . The fourth parameter is the strength of selection *s≡ α*^2^*/V*_*s*_ relative to the rate of return of population to a demographic equilibrium, *s/r*^*∗*^, and finally the fifth parameter is the scaled mutation rate, *µ/r*^*∗*^. Further details on the model and its scaling are provided in the Methods.

**Table 1.**
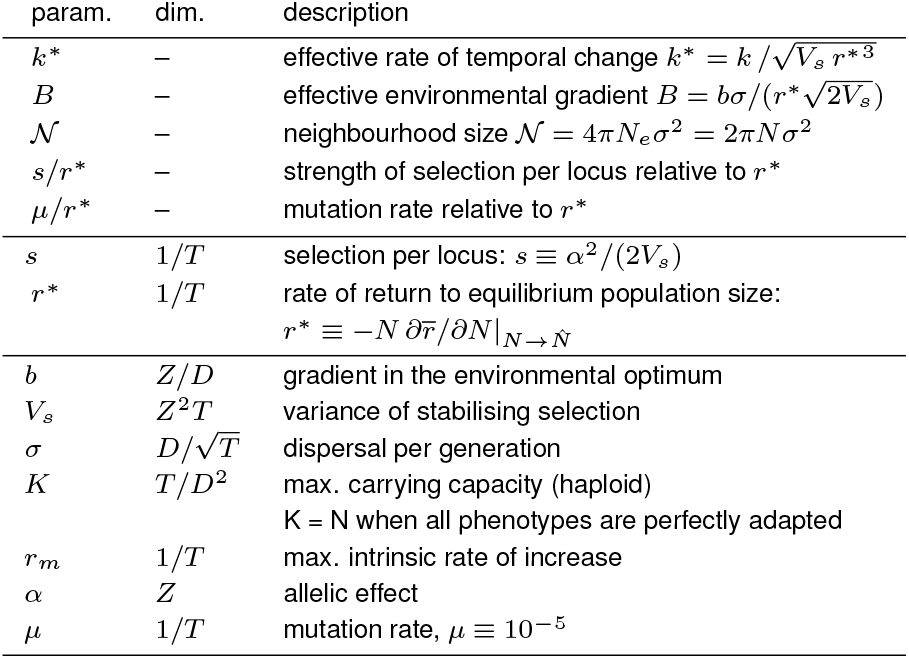
Five scale-free parameters: *B, k*^*∗*^,𝒩, *s/r*^*∗*^and *µ/r*^*∗*^ describe the system. The middle column gives the dimensions of the parameters, where T stands for time, D for distance and Z for trait.

### Known predictions: fixed variance or no temporal change

The rate of adaptation increases with the product of selection gradient and additive genetic variance (53, 54); see Eq. 3. The genetic variance itself increases with the effective environmental gradient *B* such that in the absence of stochastic and demographic effects, the scaled genetic variance is 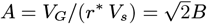 (55). We might thus expect that with increasing heterogeneity of the environment, measured by the effective environmental gradient *B*, genetic variance will steadily rise and thus support adaptation to ever-increasing temporal change *k*^*∗*^: yet, this would only be true in infinitely large populations or under soft selection. Increasing spatial heterogeneity aids adaptation until the load from increasing genetic variance becomes too large. As the proportion of population sufficiently adapted to the local environment declines, the elimination of maladapted combinations will eventually drive the population to extinction. Furthermore, long before the deterministic extinction, genetic and demographic stochasticity reshape the dynamics.

The influence of the effective environmental gradient *B* and the effective rate of temporal change *k*^*∗*^ on the rate of species’ range and niche expansion is demonstrated in Fig. 1. It shows that there is a fundamental deterministic bound to species’ range expansion at 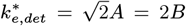 (red dashed line), which extends from the threshold derived by (32) for a given genetic variance in the absence of spatial structure. The depicted line uses the predicted genetic variance maintained by gene flow across heterogeneous environments in the absence of genetic drift. Yet, adaptation to temporal change fails considerably before this upper bound is reached. This is because genetic drift depletes genetic variance when neighbourhood size 𝒩 is small, which hinders adaptation. In the absence of temporal change, the limit where continuous adaptation to environmental gradients is possible, accounting for genetic drift, is given by 𝒩 ⪆ 6.3 *B* + 0.56 (51). This limit is derived using scaling arguments complemented by a numerical approximation. Beyond this limit, genetic drift is too strong relative to selection for stable clines to form. This threshold is depicted by a green diamond 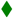 in Fig. 1.

**Fig. 1.**
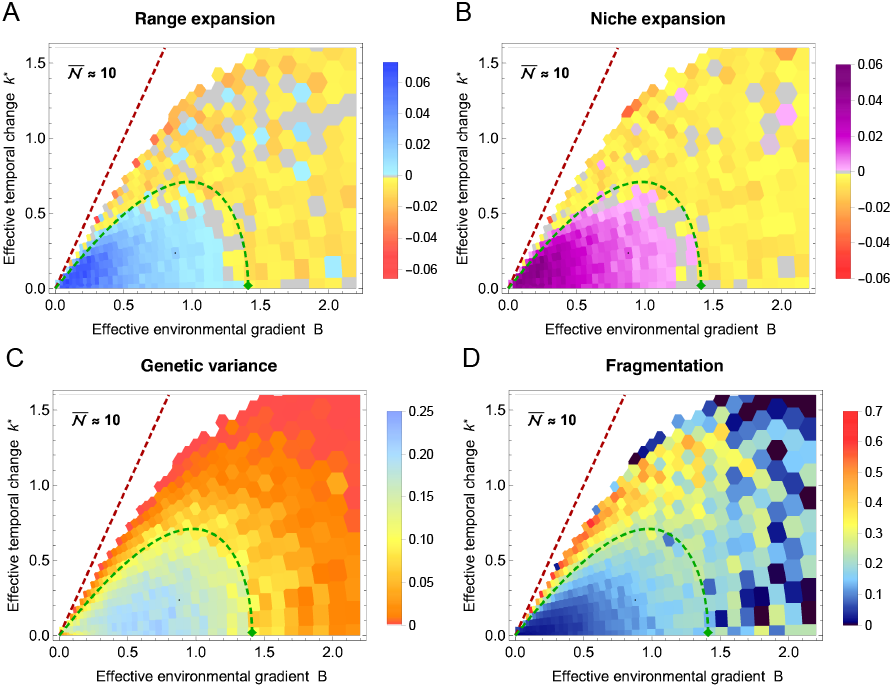
Evolution of species’ range, ecological niche and local genetic under spatial and temporal change. Species’ range (defined by the width of the occupied habitat) and niche (defined by the realized trait space) expand when sufficient variance is maintained by the gene flow across the environmental gradient and contract or fragment otherwise. Each rectangle represents an independent simulation from an array of changing environmental gradients *b* and rates of temporal change *k*. The expansion threshold is shown as a green dashed curve, approximated by 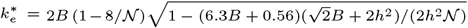 The deterministic extinction limit 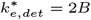 is depicted as a red dashed line. (A) The rate of range expansion relative to the initial range size (using the widest range size across the neutral dimension) is shown in blue, while the rate of niche expansion is shown in purple in (B). The rates of range and niche contraction are shown in orange and red hues. Grey indicates no significant change (95% CI) and white space represents extinctions within 500 generations. (C) Local genetic variance initially increases with both spatial and temporal variability but collapses abruptly at the expansion threshold, where the effect of genetic drift overwhelms selection. Variance is estimated as the mean local variance in the central quarter of the habitat *x*, averaged over the neutral space *y*. (D) Species’ range and niche fragment abruptly as temporal change increases. Fragmentation is measured as the proportion of unoccupied space within the species’ range. Fragmentation correlates with an increased probability of local extinction, resulting in high stochasticity in fragmentation on very steep gradients because fragmentation of the last subpopulation is zero. Median neighbourhood size 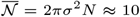 ranges between 8 and 17 (and is higher for lower *B* and *k*^*∗*^, where the overall demographic load is lower); median total population size is about 3.2 *·* 10^3^ for all surviving populations, ranging from 450 for expanding populations to 3 *·* 10^5^ at *T* = 500 generations. See Methods for parameters.

### Expansion threshold

When the optimum changes through time, the above limit to adaptation changes. A moving optimum induces lag load, which imposes a demographic cost that further reduces the neighbourhood size, thereby depleting genetic variance as genetic drift increases. In changing environments, the expansion threshold is jointly determined by the effective environmental gradient *B*, the neighbourhood size 𝒩, and the effective rate of temporal change *k*^*∗*^. Its approximate form, expressed for the critical effective rate of temporal change 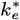, is given by 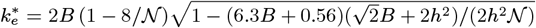, and is shown as the green curve in Fig. 1. Here *h*^2^ ≡ *V*_*G*_*/V*_*P*_ denotes the narrow-sense heritability of the additive trait. The derivation of the threshold is partly heuristic: while it utilises the analytical non-dimensionalization and approximates the effects of genetic drift on genetic variance, it neglects some interdependencies (e.g., genetic variance also evolves in response to selection imposed by the lag).

This approximate threshold marks the transition between expanding, well-adapted populations, in which variance is maintained by staggered clines in allele frequencies, and collapsing populations in which genetic drift overwhelms selection, driving erosion of clines in allele frequencies and depletion of genetic variance. Notably, it is driven by the balance between only three compound parameters *k*^*∗*^, *B*, and 𝒩. In contrast, rates of adaptation and range expansion are also influenced by the mutational input *µ/r*^*∗*^ and by the selection strength per locus, *s/r*^*∗*^. The robustness of the threshold to selection per locus *s≡ α*^2^*/V*_*s*_ is demonstrated in Fig. S3. The effect of selection per locus is negligible in two-dimensional habitats as the effect of genetic drift on the clines is hardly modulated by weak selection (e.g. in polygenic traits, where the contribution of each locus is small) (56) – it is determined by the neighbourhood size, 𝒩 ∼ *σ*^2^𝒩 . This is not the case in one-dimensional habitats, where the effect of genetic drift on the cline depends on selection, *via* 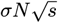 (39, 40, 57).

Genetic variance is maintained by gene flow across the environment and reduced by genetic drift. In the absence of genetic drift, the tension between local adaptation and gene flow across heterogeneous environments leads to staggered clines, with local genetic variance of 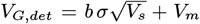 (55). Mutational variance is given by *V*_*m*_ = 2*µ n*_*l*_*V*_*s*_, where *n*_*l*_ denotes the number of loci – and in this context, it is small relative to the variance maintained by gene flow. When genetic drift increases relative to selection, these clines become increasingly rugged, shift in position, steepen, and eventually erode (39, 58, 59). Yet, the rugged allele frequencies may still contribute to adaptation even when they do not resemble a conventional cline (60, 61).

There is no exact analytical expression for the effect of genetic drift on a cline, and thus on genetic variance, in two-dimensional habitats. However, we can approximate the decrease in genetic variance due to drift as *V*_*G*_ *≈ V*_*G,det*_(1 − 8/𝒩) for 𝒩 *>* 8 – see Fig. 2. This estimate assumes that the reduction in genetic variance in two dimensions takes the same form as in one dimension (40), using the dimension-appropriate neighbourhood size, and with the constant of 8 numerically fitted. When the optimum changes in both time and space, the selection arising from the distance between the trait mean and the optimum drives the emergence of further clines in allele frequencies, which, locally, bring the trait closer to the optimum. This drives an increase in genetic variance that is stable over longer time scales relative to well-mixed populations (18) because these clines can shift through space. More generally, Fig. 2 demonstrates that neighbourhood size, rather than total (effective) population size, fundamentally drives the increase in genetic variance. Quantitative analysis of the evolution of variance in both spatially and temporally variable environments will be the subject of a forthcoming paper.

**Fig. 2.**
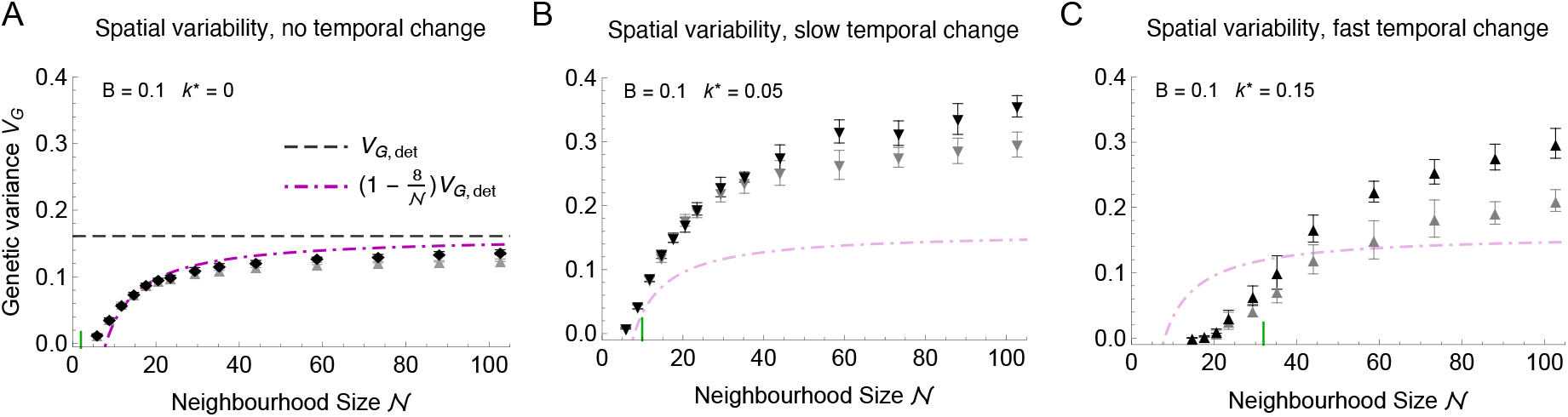
Genetic variance is significantly reduced when neighbourhood size 𝒩 is low. As the effective rate of temporal change *k*^*∗*^ increases, larger neighbourhood size is necessary so that the population can sustain itself: the expansion threshold (in terms of 𝒩 (*k*^*∗*^, *B*)), illustrated by, is roughly at the inflection point for the increase of genetic variance with 𝒩 . (A) In stable environments, the decrease in genetic variance due to drift in two-dimensional habitats approximately follows *V*_*G*_ *≈ V*_*G,det*_(1 *−* 8*/𝒩* ), shown as a dash-dotted magenta curve. Deterministic variance (dashed line) is given by 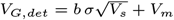 (see text). (B) When the environment changes slowly through time (at 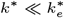), genetic variance can increase substantially as newly selected alleles spread through the population.The increase of genetic variance with *k*^*∗*^ is non-monotonic – with fast temporal change (C), increasing genetic drift suppresses the increase in variance. Black symbols: neutral habitat width is fixed at *y*_*d*_ = 100, so that total population increases with 𝒩 and the habitat remains truly two-dimensional. Gray symbols represent simulations where neutral habitat width scales inversely with carrying capacity (*y*_*d*_ = 600*/K*), ensuring similar census sizes as 𝒩 increases (*N*_*tot*_ *≈* 1.5 *·* 10^5^). When the neutral habitat width *y*_*d*_ becomes too narrow due to this scaling, genetic variance increases less than expected (gray, *y* ⪅ 50 for 𝒩 ⪆ 60). This is largely because in the limit of a one-dimensional habitat (*y* = 1), the strength of genetic drift scales with 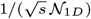 (40, 48, 50), where the one-dimensional neighbourhood size is 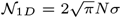 (34). Error bars show standard deviations for 10 replicates. Parameters as in Fig. 1 with *b* = 0.225, *k* = 0 (A), *k* = 0.06 *≡ k*^*∗*^ = 0.05 (B) and *k* = 0.18 *≡ k*^*∗*^ = 0.15 (C), run time *T* = 300 generations and neighbourhood size 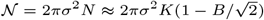, where local carrying capacity ranges from *K=*2 to *K=*35 .

### Fragmentation

As the rate of temporal change increases, a species’ range may fragment due to the feedback between genetic drift and maladaptation: see Figs. 3 and 4. Fragmentation can occur abruptly across the entire geographic range, driven by the following feedback loop: as the trait mean lags further behind the moving optimum, population size decreases due to maladaptation, and genetic drift erodes genetic variance. This, in turn, reduces the rate of adaptation, which further increases the lag load. The lag load continues to rise through this feedback loop, increasing genetic drift and reducing the population growth rate (Malthusian fitness) towards zero. Eventually, the population can no longer maintain continuous adaptation and fragments: see Movie S1.

**Fig. 3.**
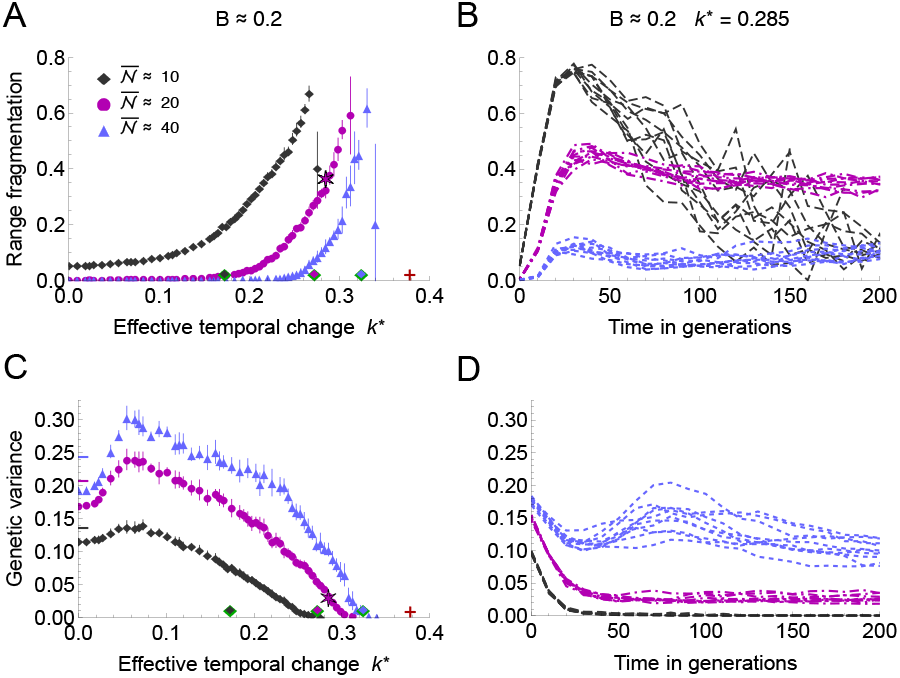
Species’ range fragments under rapid temporal change and small neighbourhood size. (A) Range fragmentation increases with the effective rate of temporal change *k*^*∗*^ and with decreasing neighbourhood size 𝒩 . The expansion threshold 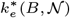 is shown as green-outlined diamonds with coloured centres matching the respective neighbourhood sizes (and it corresponds to range fragmentation higher than about 0.2). The deterministic extinction threshold 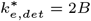 is marked with a red cross. (B) Beyond the expansion threshold, the onset of fragmentation is rapid and associated with a loss of genetic variance. The apparent decrease in fragmentation for 𝒩 ≈ 10 is due to only a few fragments remaining in fast-collapsing populations. (C) Genetic variance decreases with genetic drift (i.e., decreasing neighbourhood 𝒩size ) and is highest for intermediate rates of temporal change before declining again as genetic drift overwhelms adaptation. Short horizontal dashes at *k*^*∗*^ = 0 mark the predicted genetic variance in stable environments, accounting for genetic drift. (D) Genetic variance erodes as populations fragment when neighbourhood size is small. Only for the highest neighbourhood size 𝒩 = 40 (above the expansion threshold for *k*^*∗*^ = 0.285, *B* = 0.2), genetic variance temporarily increases again due to local selective sweeps, before the variance stabilizes at about *V*_*G*_ = 0.1. Parameters as in Fig. 1 with *b* = 0.4 (and *k* = 0.31 *≡ k*^*∗*^ = 0.285 for B, D); the magenta star marks an illustration shown in Fig. 4. Neighbourhood sizes correspond to local carrying capacities *K* = 5, 10, 20, and are slightly below the maximum attainable neighbourhood size 𝒩 = 2*πσ*^2^*K*. Error bars show standard deviations for 10 replicates. Runtime 500 generations, with first 200 generations shown in (B, D).

**Fig. 4.**
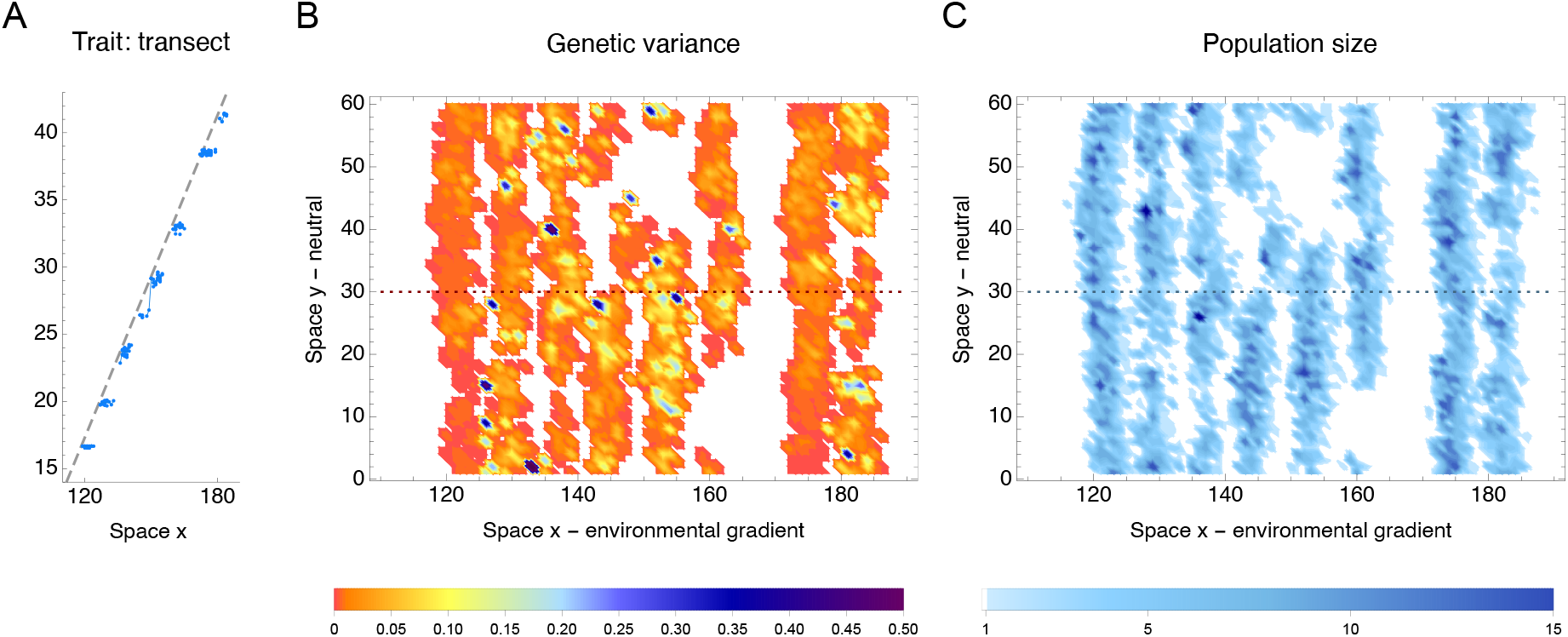
Example of species’ range fragmentation due to a fast temporal change. (A) In fragmented populations, trait mean (blue line) no longer tracks the optimum (grey dashed line) but instead, disjunct subpopulations are each adapted to a single optimum and move through space tracking their niche. Blue dots show individual trait values. The subfigure (A) shows a transect at the central neutral habitat (*y* = 30), marked by the dotted line in (B, C). (B) Genetic variance is depleted considerably below the expected value of 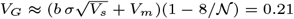, although local spikes of high variance (blue hues) persist. (C) Local population density (population size contained in a single deme) is close to the carrying capacity *K* = 10, yet 40% of the space is unoccupied, and the population gradually contracts over time. Subpopulations are connected *via* gene flow along the neutral habitat (y) but largely isolated along the environmental gradient (x). Parameters are the same as in Fig. 3 (K = 10, magenta), with *k*^*∗*^ = 0.285, with this example indicated by a magenta star there. This example is chosen just beyond the expansion threshold 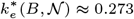, where some clines in allele frequencies still persist (see Fig. 5) and therefore genetic variance is not uniformly low. The dynamics of fragmentation is illustrated in Supplementary Movie S1, demonstrating that fragmentation occurs abruptly and follows a marked increase in lag (and thus lag load) behind the moving optimum.

The resulting metapopulation is now adapted to a series of distinct niches and may persist for some time, although its resilience is low. The lag load in the metapopulation is low, as each subpopulation tracks its old optimum through space, and they also have a low standing load (because genetic variance is depleted), as well as a low dispersal load (because the subpopulations are disjunct with little or no gene flow between them). However, fragmented populations have a very limited ability to adapt due to their depleted genetic variance. In addition, fragmented metapopulations are subject to large demographic fluctuations, which further reduce their resilience.

These two modes of adaptation to an environmental gradient were predicted by the classic phenotypic model (6) with fixed genetic variance; extended to include temporal change by (52). In the first mode, there is sufficient genetic variance, and the population can adapt uniformly to the spatial gradient. In the second mode, genetic variance is low relative to the effective spatial gradient, and the population is only adapted to a single optimum – as in the fragmented populations. Note that a species’ range can also fragment in the absence of temporal change when the neighbourhood size is small, particularly beyond the expansion threshold: see (51) and Fig. S4 (for *k*^*∗*^ = 0). Due to strong genetic drift, clines in allele frequencies become increasingly rugged, steep, and unstable. As clines steepen, genetic variance erodes and can become too low to support a continuous adaptation, *cf*. (6) and (57, Fig. 4). Demographic fluctuations, while not a primary driver, also contribute to fragmentation and accelerate the collapse of adaptation, because maladaptation increases due to higher genetic drift at low population sizes, which in turn drives even greater demographic stochasticity.

The theory predicts that an increase in dispersal may reverse fragmentation by enlarging the neighbourhood size. This effect is illustrated in Fig. 5, which depicts the dynamics of the underlying cline frequencies. Upon an increase in dispersal that raises the neighbourhood size beyond the expansion threshold (B *vs*. C), clinal adaptation gradually recovers, and the population regains continuity in both the realised trait (niche space) and geographic space.

**Fig. 5.**
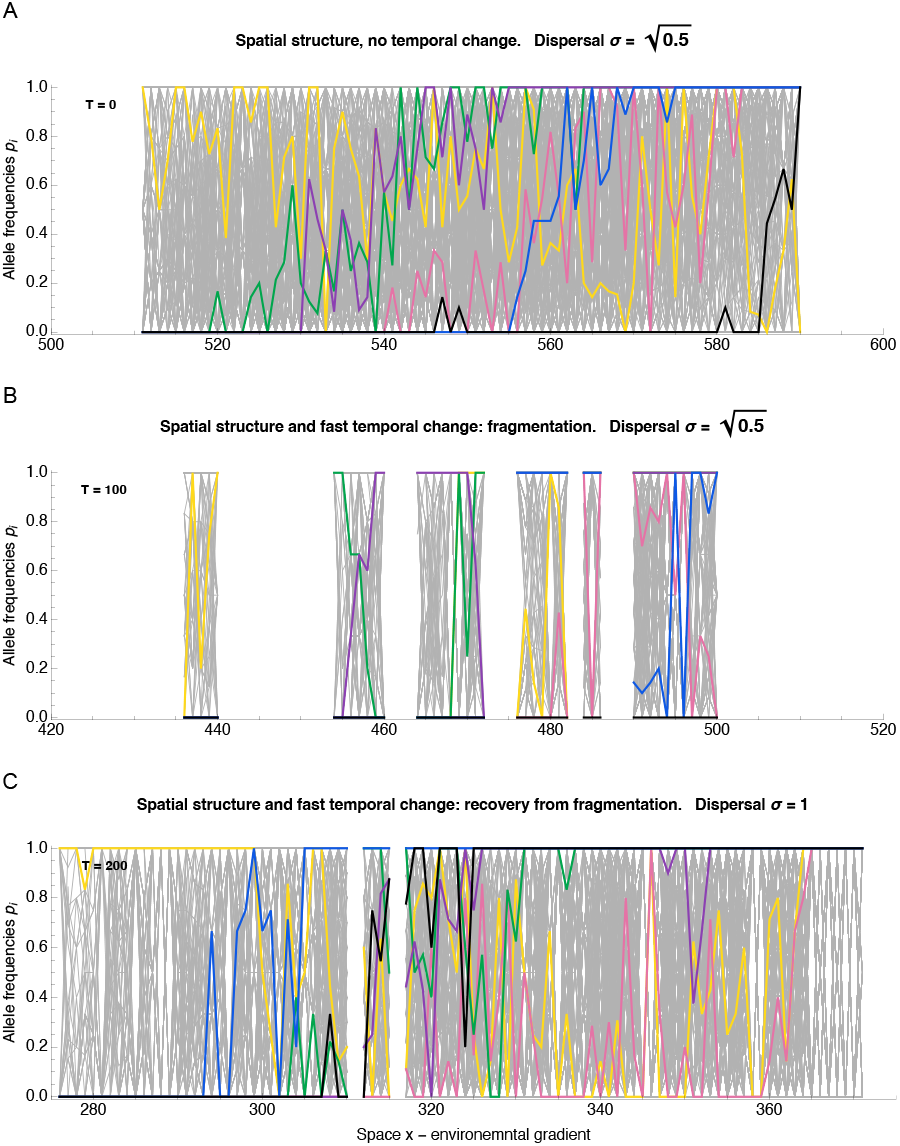
Increased dispersal restores continuous adaptation in fragmented populations. Subfigures (A–C) depict in colour six illustrative loci over a grey background showing all polymorphic loci. (A) Continuous adaptation to an environmental gradient is maintained by a series of staggered clines. These clines are rugged due to strong genetic drift, yet together they maintain a smoothly changing trait adapted to the environmental gradient. (B) Clinal variation is substantially reduced in fragmented populations. Upon a fast temporal change with an effective rate of *k*^*∗*^ = 0.285 over 100 generations (marginally beyond the expansion threshold 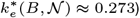, the population fragments, and most of the clinal variation is lost. (C) An increase in local dispersal *σ* enlarges the neighbourhood size and reverses fragmentation under the same rate temporal change. With a wider dispersal kernel (*σ* changes from 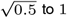 ), genetic variance maintained by dispersal across the environment increases, while genetic drift decreases. After 200 generations with broader dispersal (and thus weaker genetic drift), most of the variation is restored. Clines appear more rugged than in the absence of temporal change (A). Even at the limit of low stochasticity, cline shape is no longer necessarily monotonic, because local selective sweeps can create “bumps” in cline frequencies that bring the trait locally closer to the moving optimum. Parameters are the same as in Fig. 4. The corresponding selection per locus for illustrative clines is *s* = *α*^2^ */*(2*V*_*s*_): *s ≈* 0.03 (black), *s ≈* 0.016 (blue), *s ≈* 0.009 (purple), *s ≈* 0.007 (green), *s ≈* 0.0046 (pink), *s ≈* 1.5 *×* 10^*−*6^ (yellow). Darker colours indicate stronger selection, with the yellow cline effectively neutral. In the absence of temporal change (A), the neighbourhood size calculated from ecological parameters using the mean local deme size and parent-offspring dispersal distance *σ*, is approximately 𝒩 ≈ 25; while for the recovered population with a wider dispersal kernel (C), it is 𝒩 ≈ 31, despite local density decreasing from 8 to 5 due to elevated dispersal load. Census size drops from 39079 individuals (A) to 13079 individuals (B) and then recovers to 31767 individuals in (C).

### Dispersal and neighbourhood size

Larger neighbourhood size promotes adaptation by mitigating genetic drift and supporting adaptation to steeper environmental gradients *B* and faster temporal change *k*^*∗*^. Supplementary Figs. S4 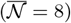 *vs*. S5 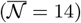 show that the expansion threshold 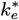 increases with neighbourhood size 𝒩, as predicted – even when the census population size is kept invariant (by scaling the neutral habitat width inversely with the local carrying capacity).

Neighbourhood size also increases with dispersal, but so does the effective environmental gradient *B*. However, in two dimensions, neighbourhood size 𝒩 = 2*πσ*^2^*N* increases with the square of dispersal distance *σ*^2^, while the effective gradient 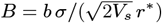 increases only linearly. Therefore, when genetic drift is strong, increasing local dispersal aids adaptation. Fig. S6 demonstrates that the benefits of reducing genetic drift *via* increased dispersal outweigh the costs associated with the increasing dispersal load: when both the effective environmental gradient *B* and the neighbourhood size 𝒩 increase with local dispersal, the drift-induced expansion threshold is approximately replaced by the deterministic limit,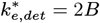.

So far, we have assumed diffusive dispersal with a Gaussian dispersal kernel. However, dispersal kernels may have heavier tails than implied by local diffusion. The effect of a leptokurtic dispersal kernel is illustrated by an example where the dispersal kernel is a mixture of two Gaussians, one with a substantially higher variance. Fig. S7 (B) shows an example where 10% of migrants disperse, on average, 10*×* further than the baseline dispersal distance of the remaining 90% of the population. Such fat-tailed dispersal (example kurtosis *κ* = 25.3) has little effect in facilitating adaptation to steep gradients, relative to the baseline with just local diffusion (Fig. S7 (A)). Yet, the fat-tailed dispersal kernel can facilitate adaptation to temporal change when the spatial gradient is shallow: this makes sense, as the increased mixing brings in further variants which may be beneficial under fast temporal change, while the cost of dispersing far across a shallow gradient is low.

Quantitative generalizations beyond diffusive dispersal are not trivial: both the effective environmental gradient *B* and the neighbourhood size 𝒩 change. Under diffusive dispersal, neighbourhood size is given by the ratio of the effective area of spread per generation, 2*πσ*^2^, to the per-generation coalescence rate, 1*/*(2*N* ), yielding 𝒩 = 4*π Nσ*^2^ (35, 56). Wright (34) defined neighbourhood size as the inverse of the identity-by-descent of nearby individuals, yielding the same 𝒩 under Gaussian dispersal. While Wright generalized the neighbourhood size to non-Gaussian dispersal kernels (62, p. 302-310), the inverse of generalized Wright’s 𝒩 may no longer robustly measure the strength of genetic drift. Under fat-tailed dispersal, the identity in state over local scales is dominated by local dispersal, and over wide scales by long-distance dispersal (35, 63). Thus, rare long-distance events have little effect locally (on the scales of cline widths in this model). Only when the kurtosis of the dispersal kernel is moderate, so that the diffusion approximation is not strongly violated, do simple generalizations of both the effective environmental gradient *B* and Wright’s neighbourhood size provide an accurate prediction: see Fig. S7 (C, D). Note that there is no simple approximate generalization of Wright’s neighbourhood size to arbitrary dispersal kernels using only their variance and kurtosis, *cf*. (43, Box 2); see S7 for details.

### Adaptive phenotypic plasticity

Last, I briefly explore the effects of adaptive phenotypic plasticity. Fig. S8 demonstrates that adaptive phenotypic plasticity does not extend the range of survivable rates of temporal change in the same manner as it does for spatial gradients. Adaptive phenotypic plasticity implies that the expressed trait *z* is closer to the optimum *θ* than expected based on its genetic basis alone, (*z− θ*) *→* (1*−* p) (*z−θ*), where p denotes adaptive plasticity. This means that the spatial gradient *B* appears shallower by a factor of 1 *−* p. While this reduces the demographic load and lowers the impact of genetic drift, it also diminishes the extent of the increase in genetic variance due to spatial variability. Thus, when genetic variance evolves in response to the effective environmental gradient, the effects of plasticity on adaptation to a steady temporal change largely cancel out. This finding contrasts with predictions assuming a fixed genetic variance in the selected trait, where the benefits of plasticity in response to variation in time and space are equivalent (7, 64).

## Discussion

There is an urgent need for predictive theory to forecast adaptive responses and population resilience in changing environments (20, 65–69). The existing ecological niche modelling (ENM) framework typically considers a multidimensional niche space that includes both biotic and abiotic components but it accounts neither for adaptive potential (70–72) nor for genetic drift, with only recent advances incorporating adaptive genetic diversity (73, 74). Adaptation typically involves changes across many loci, and predicting the trajectory of each locus is not feasible (75). In contrast, predictions based on higher-order parameters derived from fundamental quantitative genetics (based on additive genetic variance in the selected trait) are considerably more robust, particularly in controlled settings and over shorter time scales (76–81). Yet, already for predictions over time scales longer than a few generations, species’ range dynamics may change qualitatively when eco-evolutionary feedback is included: population growth depends on adaptation, the rate of which is fundamentally determined by genetic variance in the selected traits – and genetic variance itself evolves due to dispersal, selection, mutation, and genetic drift.

By explicitly integrating key eco-evolutionary processes and the feedback between them, this theory provides novel quantitative predictions concerning species’ range limits, adaptation, and resilience. It explains that a gradual increase in the rate of temporal change, as well as a decrease in neighbourhood size 𝒩, can lead to a tipping point where adaptation collapses. Neighbourhood size, the population accessible by dispersal within a generation, can decrease when dispersal decreases, reducing connectivity across space, or due to a reduction in local density. Beyond this eco-evolutionary tipping point, genetic drift becomes too strong relative to the strength of environmental change, leading to the collapse of clinal adaptation. The population may decline gradually, as the species’ range contracts from the margins, or fragment abruptly across the whole range. When the loss of fitness due to temporal change (*k*^*∗*^) is high, fragmentation can occur rapidly even when genetic drift is not particularly strong. In contrast, fragmentation due to spatial variation alone only occurs when the neighbourhood size is low (51). Since fragmentation is associated with both a local and global loss of genetic variance, recovery when conditions later improve is slower than the fragmentation.

Importantly, the predictive parameters are measurable: thus, the theory can be tested and offers applications in conservation genetics. Neighbourhood size 𝒩 can be estimated from ecological parameters of local density and dispersal as well as from population-genetic parameters – specifically, the rate of increase in neutral *F*_*ST*_ with distance (82, 83, Eq. 11). However, *F*_*ST*_ will be affected by population dynamics as well as by long-range migration: see (42) for estimates and further discussion. With diffusive dispersal, neighbourhood size is the number of individuals within a circle of radius 2*σ*: 𝒩 = 4*πσ*^2^*N* for a diploid organism, where *N* gives the local density and *σ*^2^ is the average axis-projected squared distance between parent and offspring. Generalizations beyond diffusive dispersal are discussed in section *Dispersal and neighbourhood size*: roughly speaking, rare long range dispersal can significantly facilitate adaptation to temporal change across shallow spatial gradients, but has little beneficial effect when the effective spatial gradient is steep – and with increasing frequency, will eventually swamp adaptation; see also (84) for the effect of long-distance dispersal on adaptation to spatial gradients only.

Parameters *B* and *k*^*∗*^ quantify the fitness loss (demographic load) due to spatial and temporal environmental variability. The dispersal load can be estimated from transplant experiments: moving a well-adapted population by a distance *σ* reduces fitness by *B*^2^*r*^*∗*2^. The fitness cost of a temporal change, *k*^*∗*2^*r*^*∗*^*/*2, over the characteristic time 1*/r*^*∗*^, is more challenging to infer: in some cases, past generations’ fitness can be assessed using seeds or diapausing animals, or, more commonly, by substituting space for time (68, 85, 86) in spatial transplant experiments – see also the Discussion in (52). The scale of this characteristic time, 1*/r*^*∗*^, can be estimated from the demographic dynamics, where the reported measure *D* of (87, 88) is approximately equivalent to *r*^*∗*^ (52, Appendix D).

There are three major limitations to this theory. First, it considers a single trait under stabilising selection and assumes no rigid genetic trade-off, which could further constrain adaptation (89). Second, it considers gradual spatial and temporal change in continuous environments. Habitat fragmentation driven by local environmental heterogeneity will generally hinder adaptation and the shift of the species’ range through space. While a full analysis of the effects of local habitat heterogeneity and habitat fragmentation would warrant a dedicated paper, Fig. S9 illustrates that extrinsic habitat fragmentation (30%of empty patches) has a significant negative effect, preventing adaptation to steep gradients and rapid temporal change. Selection also fluctuates through time, with extremes potentially incurring higher fitness costs than a steady change in the optimum with the same mean rate (90, 91). Third, the theory considers a single focal species. Although the environmental optimum can include both abiotic and biotic components (92), the theory neglects the coevolution of species over similar time scales (93, 94). While none of these concerns invalidate the scale-free parameters, they imply that the numerical value of the expansion threshold should be considered an approximate baseline, rather than a definitive value.

When dispersal is predominantly local, eco-evolutionary dynamics is determined over local scales. Therefore, the theory does not require spatial gradients to be linear and also applies to other forms of gradual change, such as environmental gradients gradually steepening across the species’ range (51, 95–97) or when neighbourhood size gradually decreases. As conditions deteriorate across space (and time), the expansion threshold may be exceeded locally. Then, the part of the species’ range beyond the expansion threshold is predicted to fragment or contract toward the new range margin. Because the dynamics operates over local scales and the predicted range margin is abrupt, the theory makes no claims regarding the abundant center of a species’ range (98, 99).

Conservation genetics and predictions from eco-evolutionary theory have been increasingly incorporated into conservation practice, and while there is still a notable separation of the fields (100), low or rapidly declining heterozygosity or effective population size *N*_*e*_ represents a clear cause for concern (101). Yet, the utility of any cut-off value for effective population size *N*_*e*_ remains contested (102–104). At the same time, assisted migration (44, 105, 106) is increasingly advocated as a strategy to facilitate adaptation under climate change. To date, no comprehensive predictive theory has incorporated the effects of gene flow across variable environments on local polygenic adaptation and species’ range dynamics under temporal change. This theory provides a framework for predicting when an increase in local dispersal, for example via assisted migration, can aid adaptation and identifies a critical tipping point beyond which populations may be at risk of collapse or abrupt fragmentation.

## Materials and Methods

The model describes the evolution of a species’ range in two spatial dimensions. Crucially, population dynamics and evolutionary dynamics are coupled through the mean (Malthusian) fitness 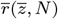,which gives the growth rate of the population, decreasing with population density and maladaptation: 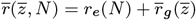.

Building on classic models of species’ range evolution (5, 6, 55), we first describe the joint model of population dynamics and the evolution of the trait mean and its variance, using stochastic differential equations. This allows us to derive the effective parameters (see Table 1), which is essential for understanding the dynamics. In the last part of the Methods, we describe an individual-based model with a polygenic trait, which is used to test the robustness of the predictions based on the effective dimensionless parameters.

### Population dynamics

The ecological component *r*_*e*_(*N* ) of the population growth rate 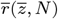 can take various forms. We assume logistic regulation, so that fitness declines linearly with local population density *N* : *r*_*e*_(*N* ) = *r*_*m*_(1 *− N/K*), where *r*_*m*_ is the maximum per capita growth rate (intrinsic rate of increase) in the limit of *N →* 0. Throughout, we drop the spatial coordinates (*x, y*) from the notation for simplicity: *N≡ N* (*x, y*). The maximum carrying capacity *K* is assumed to be uniform across space but both the growth rate and the attainable population density 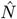decline with maladaptation.

The reduction in growth rate 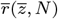 due to maladaptation is represented by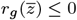. Selection is stabilising around the optimum *θ* = *b x−kt*, which changes smoothly (with a gradient *b*) along one spatial dimension (*x*), and through time *t* at a constant rate *k*. For any individual with a trait value of *z*, the drop in fitness due to maladaptation is *r*_*g*_ (*z*) = *−* (*z− θ*)^2^*/*(2*V*_*s*_), where 1*/V*_*s*_ gives the strength of stabilising selection. A population with a mean phenotype 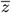 and Gaussian phenotypic variance *V*_*P*_ has its fitness reduced by 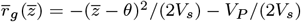. Altogether, the growth rate of the mean phenotype *z* at local density *N* is

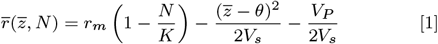

It follows that the maximum attainable density 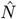 for a well adapted population, with trait mean 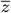 matching the optimum *θ*, is 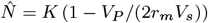.

Phenotypic variance is *V*_*P*_ = *V*_*G*_ + *V*_*E*_. The loss of fitness due to environmental variance *V*_*E*_ can, however, be incorporated into an effective intrinsic rate of increase 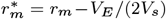. Substituting 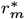 for *r*_*m*_ renders the environmental variance *V*_*E*_ a redundant parameter. At low densities, when the intrinsic rate of increase drives the population dynamics, the extra term *−V*_*E*_*/*(2*V*_*s*_) *N/K* vanishes. Accordingly, we set *V*_*P*_ = *V*_*G*_ throughout.

### Evolution of trait mean with joint population dynamics; unscaled

Population dynamics in two dimensions can be described by diffusive migration corresponding to independent Brownian motion of individuals and a deterministic growth component given by the mean Malthusian fitness 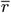, and stochastic demographic fluctuations. Random fluctuations in realized fitness are described by a demo-graphic noise term 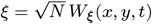, where *W*_*ξ*_ denotes Gaussian spatiotemporal white noise (107). This approximation is consistent with Poisson-distributed offspring numbers in a discrete-time model, where both the mean and variance of offspring per generation are proportional to *N* .

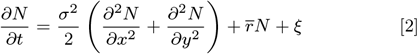

For any given additive genetic variance *V*_*G*_ (assuming a Gaussian distribution of breeding values), the change in the trait mean 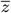 over time satisfies:

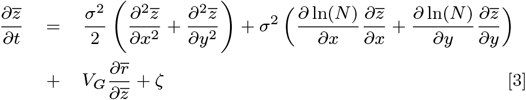

The first term gives the change in the trait mean due to diffusive migration, with a mean displacement of *σ*; the second term describes the effect of the asymmetric flow from nearby areas of higher density; the third term gives the change due to selection, given by the product of genetic variance and gradient in mean fitness (5, Eq. 2). The last term, *ζ*, gives fluctuations in the genetic variance due to genetic drift: 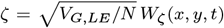, where *W*_*∗*_ represents white noise in space and time (39, 56).

Because the probability that a given individual is the parent of a particular offspring is small, Poisson distribution of the offspring number corresponds well to the binomial distribution of Wright-Fisher model (1, 3), which can be passed to the diffusion limit (48, 108). Yet, the diffusion approximation breaks down over very local scales in two dimensions because in a continuous two-dimensional habitat, two independent Brownian motions will, with probability one, never occupy the exact same place at the same time (and thus, there would be no coalescence). The individual-based simulation avoids this problem by considering a discrete lattice; a formally fully consistent analytical approach would require the introduction of a local scale of population interactions (56).

Substituting population genetic formulae for the trait mean and genetic variance into Eq. 3 gives the dynamics of allele frequencies *p*_*i*_. For a haploid model, the trait mean is 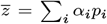 where *p*_*i*_ is the frequency of the allele with effect *α*_*i*_ at locus *i*, and *q*_*i*_ = 1 *−p*_*i*_; the genetic variance, assuming linkage equilibrium (LE), is 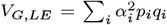. The allele frequencies (for both haploid and diploid models) then change through time due to dispersal, selection, mutation, and genetic drift, as follows:

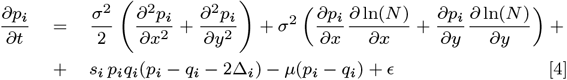

The term *s*_*i*_ *p*_*i*_*q*_*i*_(*p*_*i*_ *− q*_*i*_ *−* 2Δ_*i*_) represents the product of the heterozygosity, *p*_*i*_*q*_*i*_, and the gradient in mean fitness, 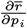. It gives the rate of change due to selection, where selection at locus *i* is 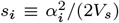, and the scaled deviation of the trait mean from the optimum is 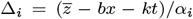 (55, Appendix 3). The fourth term describes the change due to symmetric mutation at rate *µ*, and the last term, *ϵ*, describes genetic drift (56, Eq. 7): 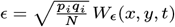, where *N* is the haploid population density.

It can be shown that in the absence of genetic drift, allele frequencies are approximately at linkage equilibrium (LE): positive linkage disequilibrium (LD) generated by dispersal approximately cancels out with the negative linkage disequilibrium generated by stabilizing selection, to first-order in the pairwise LD (57, Supplementary Information), extending from (109). Genetic drift, however, would generate positive linkage disequilibrium. At the infinitesimal limit (110), each allele has such a small effect that changes at each locus are transient, and selection operates by changing LD between loci. In general, therefore, linkage disequilibrium may develop, particularly when genetic drift is strong: we will assess the evolution of LD in a separate paper concerned with the details of the evolution of genetic variance and genetic architecture.

In the absence of genetic drift and any temporal change, genetic variance is largely maintained by gene flow, and the contribution of mutation is relatively small. Generally, the variance maintained by mutation-selection balance in a well-mixed population, *V*_*G,µ/s*_ = 2*µV*_*s*_*n*_*l*_ (where *µ* is the mutation rate per locus and *n*_*l*_ the number of loci), is expected to be considerably smaller than variance generated by gene flow across environments,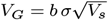 (55, 61, 111, 112).

### Rescaling and formalising in terms of allele frequencies, with evolution of variance

Following (6, 55), the model (Eqs. 2 and 3, 4) can be simplified by rescaling time *t* with the characteristic rate *r*^*∗*^ of return to the equilibrium population size, distances *x* and *y* relative to dispersal *σ*, trait *z* relative to the strength of stabilising selection 1*/*(2*V*_*s*_), and local population density *N* relative to equilibrium population density under perfect adaptation:

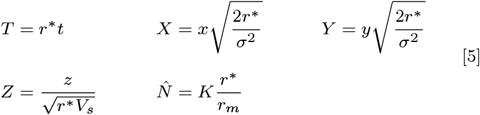

The characteristic time *T* scales with the rate of return to the equilibrium population density, defined as its negative elasticity: 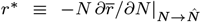; see also (87,_*∗*_ 113). When the trait mean matches the optimum, we get 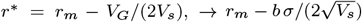. In addition, we normalise density as 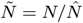, such that near the equilibrium of a well-adapted population,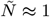. White noise is rescaled as 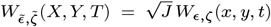,where *J* = *dx dy dt/dXdY dT* = *σ*^2^*/*(2*r*^*∗*2^); see also (39).

The rescaled equations for evolution of allele frequencies and for demographic dynamics are:

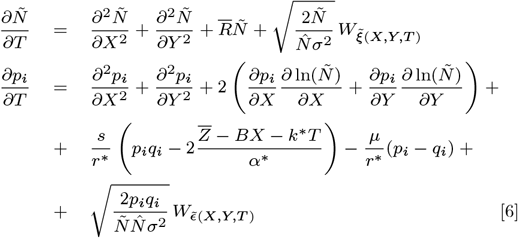

where 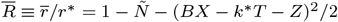.

The rescaled equations show that five dimensionless parameters fully describe the system formalized by the SPDEs. The first parameter is the effective rate of temporal change, 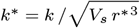. The second parameter is the effective environmental gradient, 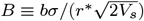. The third parameter represents the strength of genetic drift, 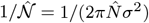, where 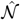 gives the neighbourhood size of a well-adapted population at equilibrium. The fourth parameter gives the strength of selection relative to the strength of density dependence, *s/r*^*∗*^; the scaled effect of a single substitution, *α*^*∗*^, also scales with *s/r*^*∗*^: 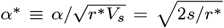The effect of this fourth parameter, *s/r*^*∗*^, is expected to be small because typically, *s* ⋘ *r*^*∗*^. The fifth parameter, *µ/r*^*∗*^, is small and can be neglected for the purpose of finding threshold conditions for range expansion *vs*. collapse. Throughout, we set the mutation rate to *µ* = 10^*−*5^.

### Individual-based simulations

Individual-based simulations in discrete time and space are used to test the robustness of the predictions based on the SPDE model with continuous time and space. The life cycle is selection → mutation → recombination → birth → migration. Generations are non-overlapping, and selfing is allowed at no cost. Every generation, each individual mates with a partner drawn from the same deme, with probability proportional to its fitness, to produce a number of offspring drawn from a Poisson distribution with mean exp[*r*(*z, N* )], where *r*(*z, N* ) = *r*_*m*_(1 *− N/K*) *−* (*z − θ*)^2^*/*(2*V*_*s*_).

The population density *N* gives the number of individuals within a single deme on the spatial lattice, *K* is the local carrying capacity, and *r*_*m*_ is the maximum intrinsic rate of increase. *θ* = *bx kt* is the optimum for the trait *z* under Gaussian stabilising selection with variance *V*_*s*_, changing linearly through space *x* with gradient *b*, and through time *t* at rate *k*. Assuming a Poisson distribution of offspring number, local population density equals the local effective population density (with the same ploidy); thus, in a haploid model with local density *N*, we have *N*_*e*_ = 2*N*, where *N*_*e*_ denotes the diploid effective population size. The unscaled parameters used in the IBM are described in the bottom section of Table 1, where the top section gives the derived dimensionless parameters of the approximated SPDE model.

The trait *z* is determined by *n*_*l*_ = 1000 diallelic loci with additive effects *α*_*i*_. These effects are exponentially distributed with mean *α* = 0.07, unless stated otherwise; this gives the mean strength of selection per locus *s* = *α*^2^*/*(2*V*_*s*_) *≈* 0.005. The genome is haploid with unlinked loci, so the probability of recombination between any two loci is 1*/*2. Mutation is symmetric and mutation rate is uniform across loci, with *µ* = 10^*−*5^.

Individuals migrate according to a discretised Gaussian distribution, approximating diffusive migration. Because the IBM runs on a two-dimensional lattice the tails of the dispersal kernel need to be truncated. This truncation is set to three standard deviations of the dispersal kernel, 3*σ*, throughout, and dispersal probabilities are adjusted so that the discretised dispersal kernel sums to 1, while ensuring the correct variance (see (114, p. 1209)). For comparison, truncation at 2*σ* would correspond to nearest-neighbour migration, where the probability of staying in place is 1*/*4, the probability of migrating to perpendicular neighbours is 1*/*8, and to diagonal neighbours is 1*/*16.

Simulations were written in Mathematica (Wolfram) and executed on the Vienna Scientific Cluster (VSC); the code will be available on Dryad upon acceptance.

### Parameters

The environmental gradient *b* and the rate of temporal change *k* are typically sampled from a range, with displayed values of *b* ranging from 0.05 to at most 2.4, and *k* ranging from 0 to at most 1.3. Gaussian dispersal per generation has a parent-offspring variance of *σ*^2^ = 0.5 (except for Figs. S6 and S7), the width of stabilising selection is *V*_*s*_ = 1, maximum growth rate *r*_*m*_ = 1.2, mutation rate *µ* = 10^*−*5^, *n*_*l*_ = 1000 loci with exponentially distributed effects with mean 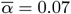 (except for Fig. S3). Initially, one third of all loci form clines that contribute to adaptation to the environmental gradient, such that within the central third of the range, the trait mean matches the environmental optimum. The allele frequencies of the remaining two thirds of loci are initially spatially uniform, with half of these loci fixed at 0 and the other half fixed at 1. These loci therefore contribute to the trait value but not to the spatial gradient in trait mean at time zero. There is no explicit genetic map, so the genomic (vector) positions of the preadapted (clinal) loci are irrelevant. The space is represented by a two-dimensional lattice *x*_*d*_ *× y*_*d*_ with spacing between demes *δx* = 1, and is wide enough so that the species’ range does not reach the edges along the habit dimension with the environmental gradient, *x*_*d*_ *≥* 600 in the course of the simulation, and that the stochastic fluctuations of allele frequencies occur in truly two-dimensional habitat: the number of demes along the neutral dimension is *y*_*d*_ = 600*/K* = 100. Burn-in time is 200 generations with no temporal change in the optimum; run time is 500 generations.

## Acknowledgements

I would like to thank Nick Barton, Roger Butlin and Richard Nichols for discussions and detailed comments on earlier drafts, and Stuart Baird, Andrea Betancourt, Puneeth Deraje, Louise Fouqueau, Jan Hrček, Patrik Nosil, Pavel Payne and Jason Sexton for their comments on the manuscript.

## Supplementary Materials

Supplementary Figures S1 to S9

Supplementary Movie S1

