## Supplementary Figures and Legends for "Evolution of Species’ Range and Niche in Changing Environments"

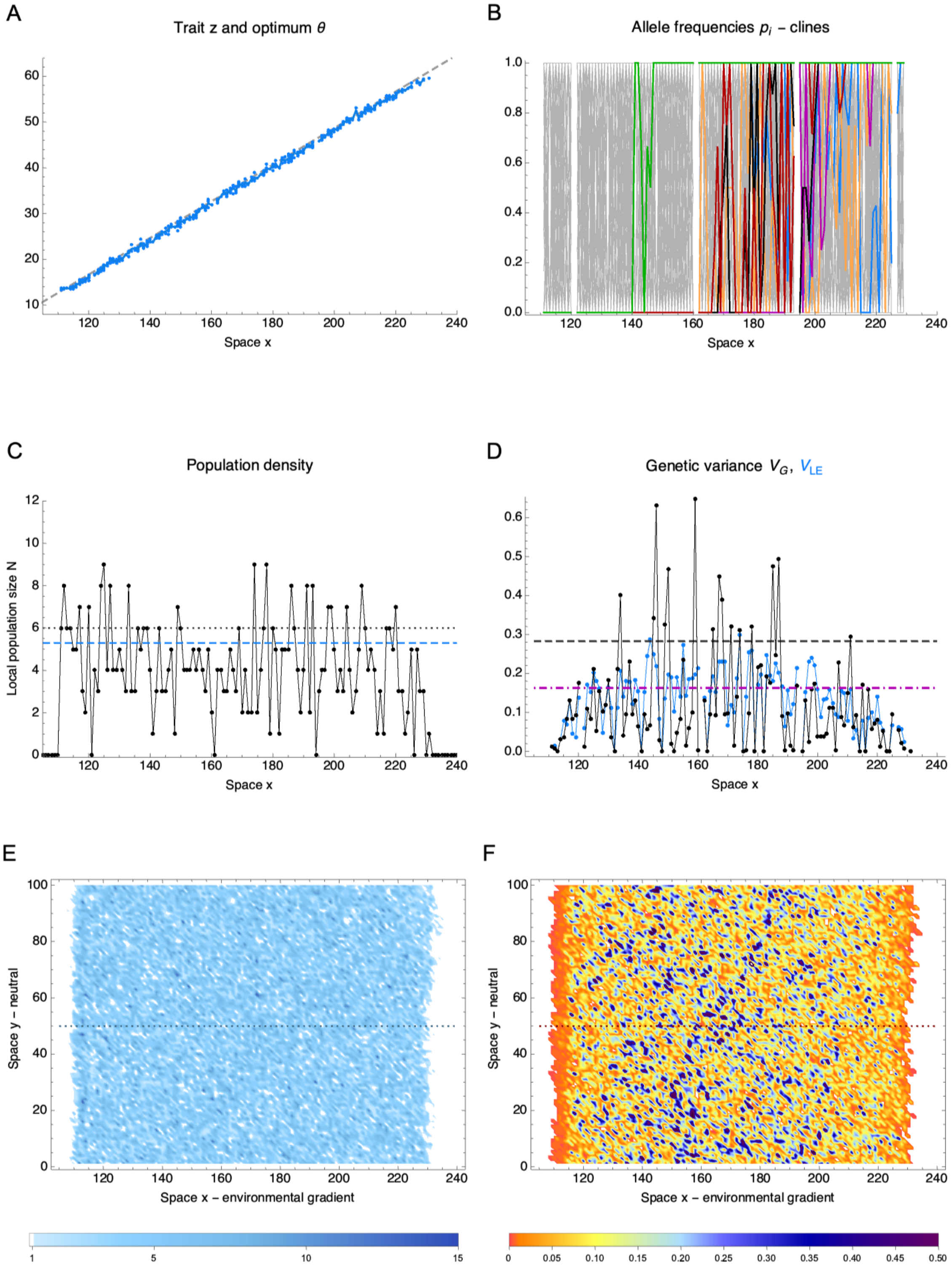


**Fig. S1 Illustration of coupled evolutionary and population dynamics with slow temporal change $k^* = 0.1$ and effective spatial gradient $B = 0.2$.** (A–D) a transect through the central neutral habitat; (E, F) a two-dimensional view across space. (A) As the optimum changes slowly through time (to the left), trait mean $\bar{z}$ (blue; line: mean, dots: individuals) lags just slightly behind the optimum $\theta$ (gray dashed line). (B) Clines which underlie the trait are nevertheless very noisy: this is due to the combination of genetic drift and temporal change. All clines are shown as a grey background, six coloured clines illustrate the observed cline shapes. (C) Local population size is on average close to the deterministic expectation in the absence of temporal change (blue dashed line); the dotted line gives the carrying capacity $K = 6$. (D) Genetic variance is even more noisy, yet the average is not far from the expected value in the presence of drift (but no temporal change), shown in dash-dotted magenta (see Fig. 2.) Dashed black line gives the expected variance in the absence of genetic drift and temporal change ($V_G = b\sigma\sqrt{V_s}$). Total genetic variance is shown in black, with its linkage equilibrium component in blue: while overall they are close, there may be significant departures from linkage equilibrium locally. (E) The population is continuous, while local population size is around the expected value. (F) Genetic variance varies across space with an average around the expected value away from the (variation-depleted) edges. Parameters as in Fig 3, with $k^* = 0.1$. Mean neighbourhood size $\mathcal{N} = 14$, total population size $N_{tot} = 5 \cdot 10^4$.

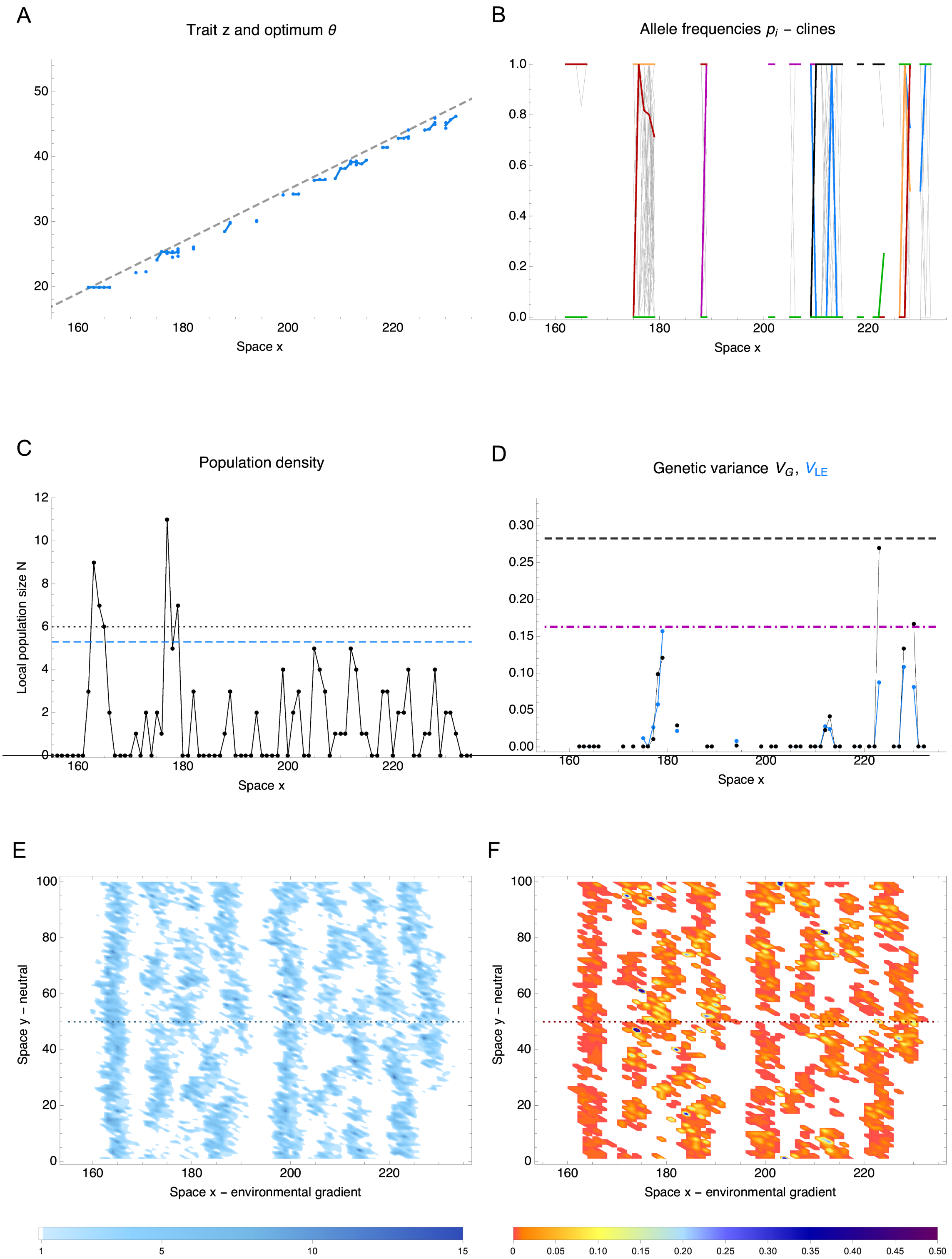


**Fig. S2 Illustration of coupled evolutionary and population dynamics with fragmentation under fast temporal change $k^* = 0.26$ and effective spatial gradient $B = 0.2$.** (A–D) a transect through the central neutral habitat; (E, F) a two-dimensional view across space. (A) As the optimum changes through time (to the left), trait mean $\overline{z}$ (blue; line: mean, dots: individuals) lags behind the optimum $\theta$ (gray dashed line). (B) Clines which underlie the trait are very noisy: this is due to the combination of high stochasticity under fast temporal change. All clines are shown as a grey background, six coloured clines illustrate the observed cline shapes. (C) Local population size fluctuates substantially, and as is on average considerably lower than the deterministic expectation in the absence of temporal change (blue dashed line); the dotted line gives the carrying capacity $K = 6$. (D) Genetic variance is depleted yet in some patches it is close to the expected value in the presence of drift (but no temporal change). The expected value is shown in dash-dotted magenta; see Fig. 2 for more details. The dashed black line gives the expected variance in the absence of genetic drift and temporal change ($V_G = b\sigma\sqrt{V_s}$). Total genetic variance is shown in black, with its linkage equilibrium component in blue: while overall they are close, there may be significant departures from linkage equilibrium locally. (E) The population is highly fragmented yet still connected along the neutral habitat (y). (F) Genetic variance is mainly depleted (red), maintained only in a few small patches. Parameters as in Fig 3, with $k^* = 0.26$ above the expansion threshold $k^*_e(\mathcal{N}, B) = 0.2$. Mean neighbourhood size $\mathcal{N} = 10$, total population size $N_{tot} = 1.2 \cdot 10^4$.

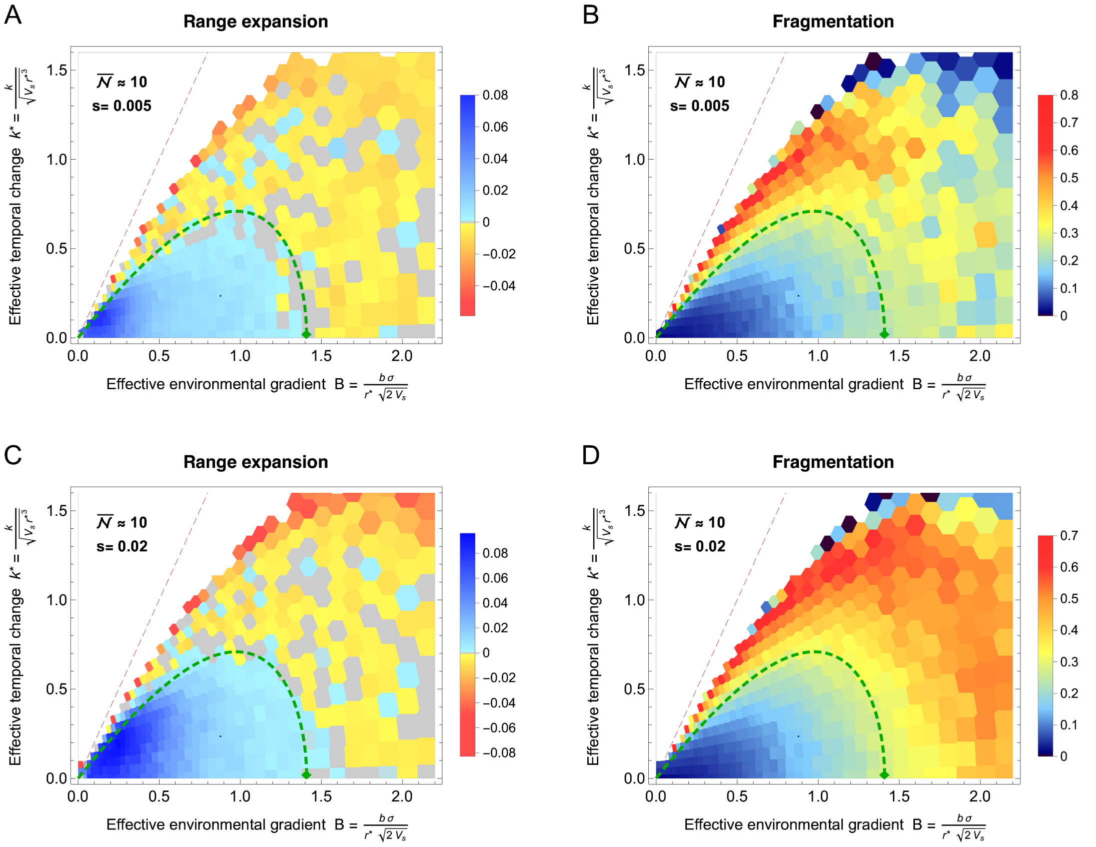


**Fig. S3 The expansion threshold (dashed green curve), which describes the tension between polygenic adaptation and genetic drift, is approximately independent of the strength of selection per locus, $s \equiv \alpha^2/(2V_s)$.** However, there is a tendency for selection to overcome genetic drift marginally sooner when selection per locus is stronger ((A, B) $s = 0.005$ *vs.* (C, D) $s = 0.02$). Notice that beyond the expansion threshold, fragmentation is more extensive (and the signal is less noisy) under stronger selection per locus (D), despite a slightly higher total population size. The dashed red line gives the deterministic extinction threshold. Parameters are as in Fig. 1 with allelic effects $\alpha$ fixed at $\alpha = 0.1$ for (A, B) and $\alpha = 0.2$ for (C, D); the number of loci scales inversely with $\alpha$ to 1000 and 500, respectively, thereby ensuring comparable range sizes at time zero.

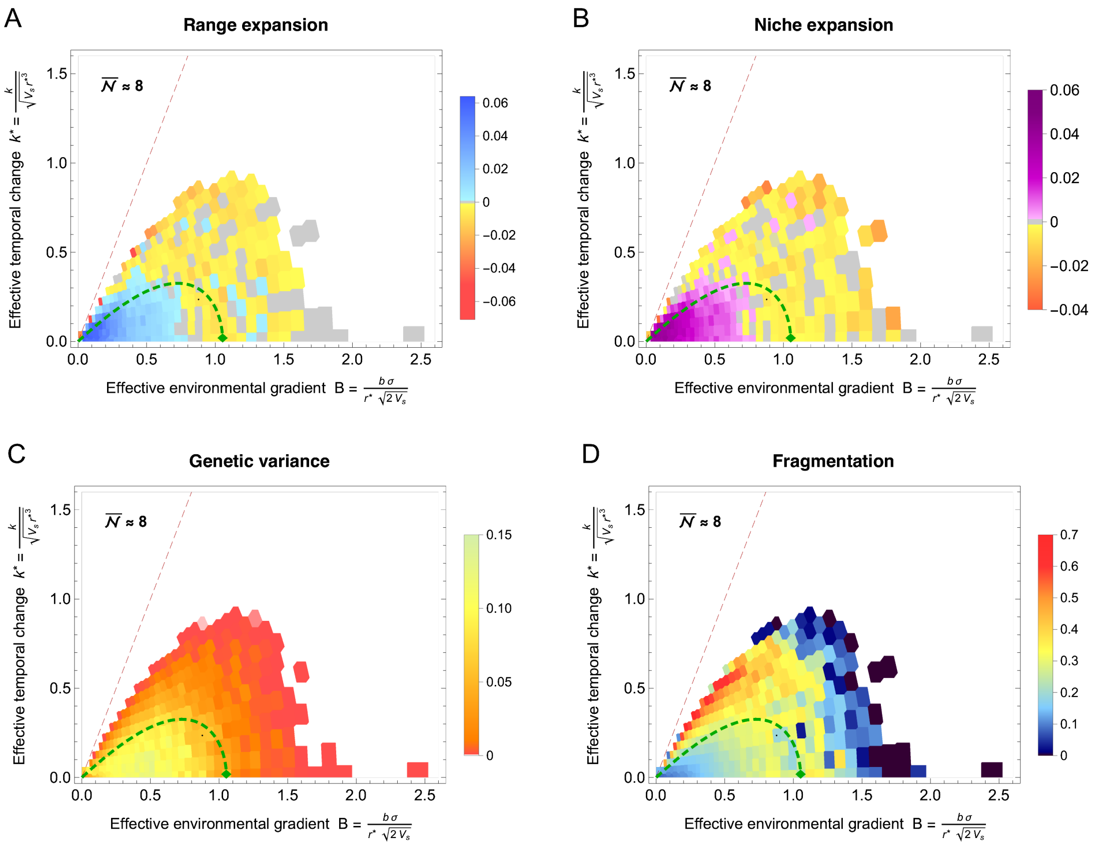


**Fig. S4 Coevolution of species' range, niche, and local genetic variance: low neighbourhood size.** As neighbourhood size decreases, expansion threshold (dashed green curve) moves toward weaker gradients $B$ and slower rates of temporal change $k^*$. It is apparent that with very low neighbourhood sizes, demographic stochasticity becomes significant, and range expansion for high effective spatial gradient $B$ ceases a little earlier than predicted. There is also a marked increase of fragmentation as $B$ increases: the subsequent decrease in fragmentation for very steep gradients means that there is only a single surviving (but collapsing) population. For weak gradients, we see that the population expands slightly beyond the approximate threshold: this is because when neighbourhood size is very low, the approximation of reduction of genetic variance starts to diverge (the prediction gives lower genetic variance than observed): see Fig. 2. Dashed red line gives the deterministic extinction threshold. Parameters as in Fig. 1 but with maximum local carrying capacity set to $K = 4$, and the neutral habitat width scaled to $y_d = 600/4 = 150$. This gives neighbourhood size with median $\overline{\mathcal{N}} = 2\pi\sigma^2 N \approx 8$, ranging from $\mathcal{N} = 6$ to $\mathcal{N} = 11$, and total population size with median $2.8 \cdot 10^3$ for surviving populations, ranging from 130 to $3 \cdot 10^5$ for expanding populations at $T = 500$ generations.

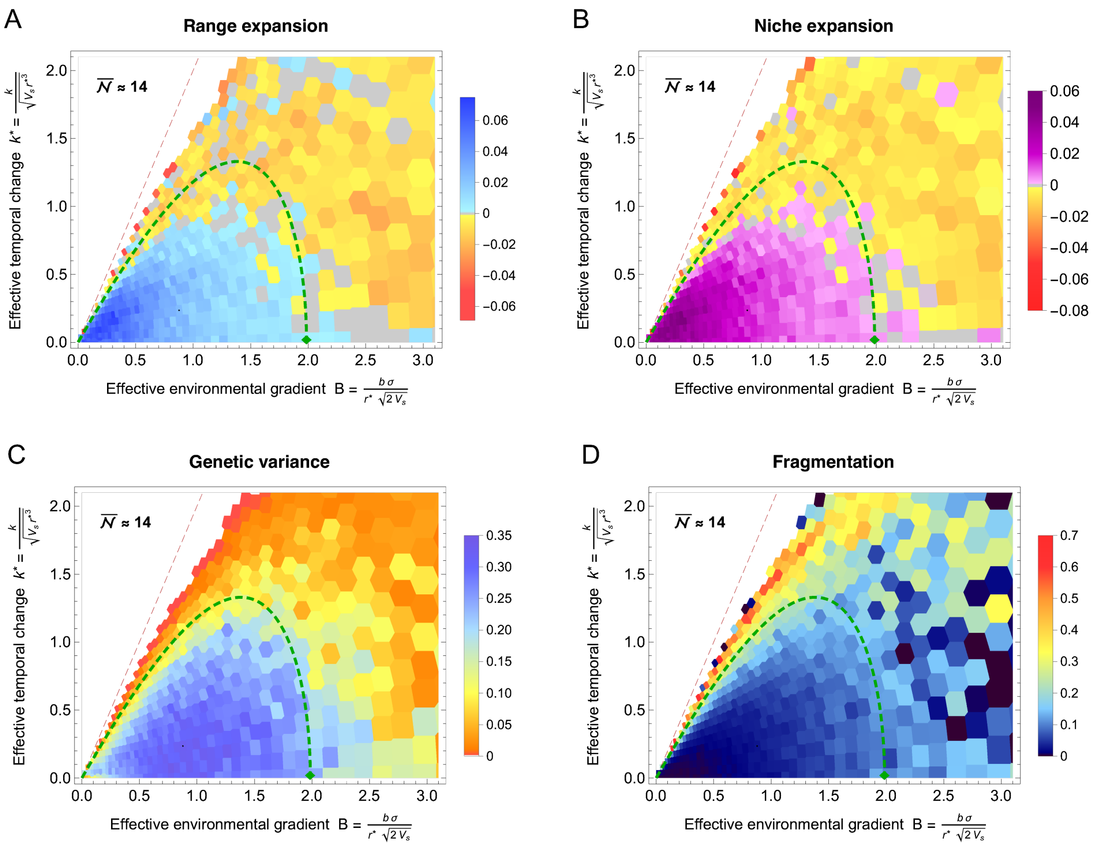


**Fig. S5 Coevolution of species' range, niche, and local genetic variance: larger neighbourhood size.** As neighbourhood size increases, species' range and niche expand for steeper gradients $B$ and faster rates of temporal change $k^*$. Under no temporal change, $k^* = 0$, the positive rate of range and niche expansion extends slightly beyond the approximate expansion threshold (dashed green curve), although the rates of expansion are very slow. In contrast, we see only a few singular, weakly expanding populations for $k^* > 0$ (some noise is expected with the CI of 95% for over 1000 independent simulations). The dashed red line gives the deterministic extinction threshold. Parameters as in Fig. 1 but with maximum local carrying capacity set to $K = 10$, and the neutral habitat width scaled to $y_d = 600/10 = 60$. This gives a neighbourhood size with median $\overline{\mathcal{N}} = 2\pi\sigma^2 N \approx 14$, ranging from $\mathcal{N} = 10$ to $\mathcal{N} = 30$ and a total population with median $3.7 \cdot 10^3$ for surviving populations, ranging from 600 to $4 \cdot 10^5$ for expanding populations at $T = 500$ generations.

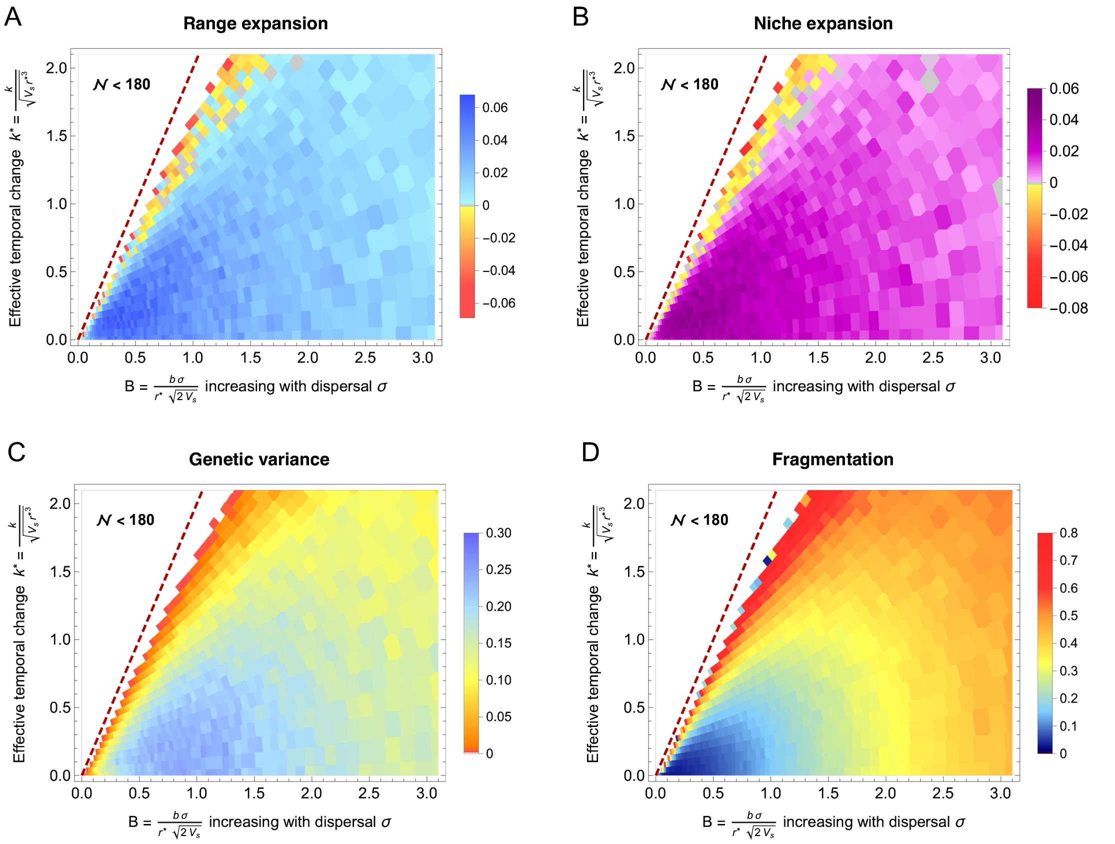


**Fig. S6 Gene flow across heterogenous environments facilitates adaptation.** This figure illustrates that when local population density is small, the benefits of reducing genetic drift by increasing dispersal $\sigma$ outweigh the costs incurred by elevated dispersal load. In contrast to Fig. 1, where $B$ increased with the spatial environmental gradient $b$, here the effective environmental gradient $B = b\,\sigma/(\sqrt{2V_s}\,r^*)$ increases with dispersal $\sigma$. This also elevates the neighbourhood size $\mathcal{N} = 2\pi\,\sigma^2\,N$, where $N$ is the population size within a deme. Then, species' range (A) and niche (B) keep expanding for increasing effective environmental gradient $B$ because the neighbourhood size $\mathcal{N}$ also keeps increasing with dispersal $\sigma$: genetic drift thus never overpowers adaptation. Increasing effective environmental gradient $B$ facilitates adaptation to temporal change $k^*$; the deterministic extinction threshold due to high lag load, $k^*_{e,det} = 2B$ is given by the dashed red line [32]. Dispersal still carries a gradually increasing demographic cost: this manifests both in a gradual decline in genetic variance (C) once the effective environmental gradients $B$ becomes very steep, and in increasing fragmentation as the effective environmental gradient $B$ increases with dispersal $\sigma$ (D). This is expected: eventually, the combined demographic loads (dispersal, variance and lag loads) will become too large and the population approaches deterministic extinction under logistic population regulation. Parameters as in Fig. 1 but with $B$ increasing with dispersal distance $\sigma$, ranging from 0.05 to 3.25, which corresponds to neighbourhood size $\mathcal{N}$ ranging from 0.1 to 180 (median $\mathcal{N} = 74$), while the local carrying capacity stays at $K = 6$. The median total population was around $1.25 \cdot 10^4$ for surviving populations, ranging from $2.2 \cdot 10^3$ to $4.4 \cdot 10^5$ for expanding populations at $T = 500$ generations.

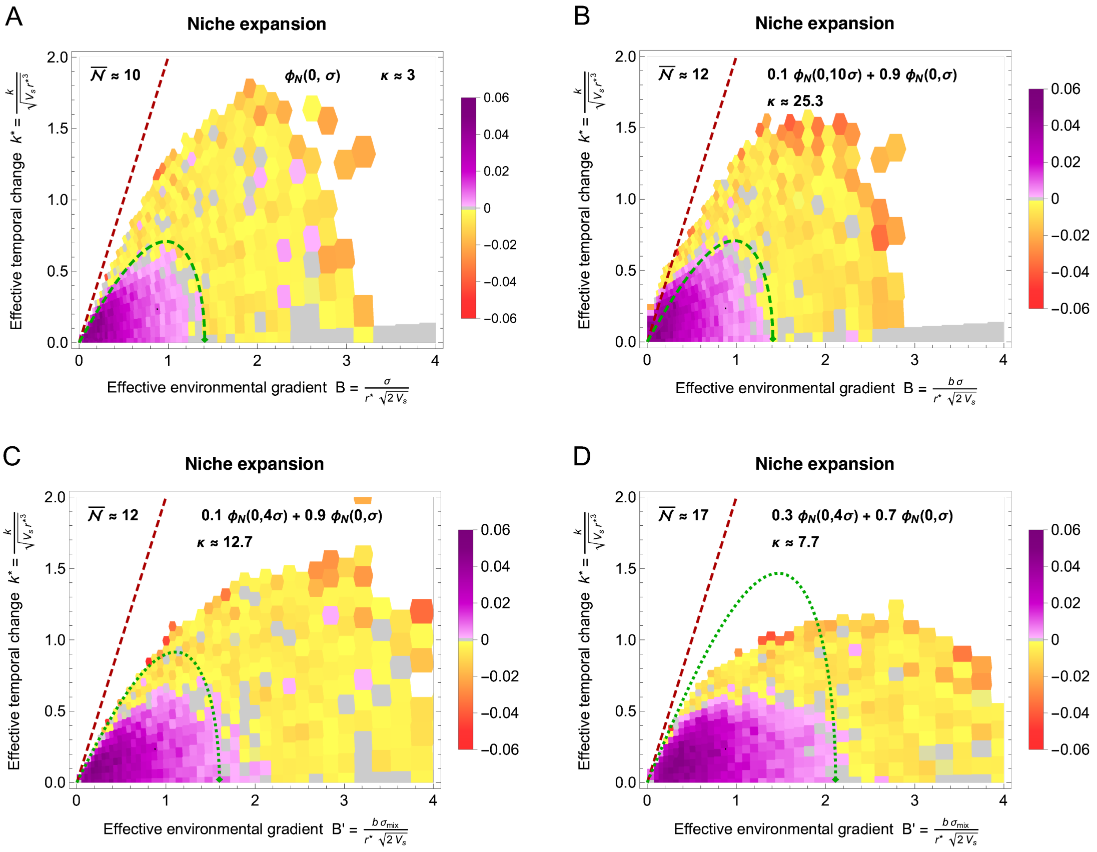


**Fig. S7 Limits to niche expansion with leptokurtic dispersal; kurtosis given by $\kappa$.** (A) shows the baseline Gaussian dispersal for comparison, with local dispersal $\sigma_1 = \sigma = \sqrt{1/2}$. (B) Long-distance dispersal can facilitate adaptation to faster temporal change on shallow to moderate spatial gradients ($B \lesssim 1$) but does not significantly facilitate adaptation to steep spatial gradients. The illustration shows a leptokurtic dispersal kernel with kurtosis $\kappa \approx 25$, generated by a mixture of two Gaussian distributions, where 10% of the population has a 10× wider dispersal kernel: $\sigma_2 = 10\sigma$. (C, D): For moderate kurtosis, the effective environmental gradient $B$ can be approximated using $\phi_{\mathcal{N}}(0, \sigma_{\text{mix}})$ as $B' = \frac{b\,\sigma_{\text{mix}}}{\sqrt{2V_s}\, r^*}$, where $\sigma_{\text{mix}} = \sqrt{a\,\sigma_2 + (1-a)\,\sigma_1}$. This stretches the experienced effective environmental gradient: due to the wider dispersal kernel, critical $B$ appears steeper though the the limiting extrinsic gradient $b$ does not change. We can also adjust the neighbourhood size using the mixture of dispersal kernels. Following Wright [62, p. 302-310], we calculate the neighbourhood size using the probability that two nearby genes share ancestry in the previous generation: $P_{id} = \frac{4\sigma_l^2}{n} \int_{-\infty}^{\infty} \int_{-\infty}^{\infty} \left( \frac{1-a}{2\pi\sigma_1^2} \exp\left(-\frac{x^2+y^2}{2\sigma_1^2}\right) + \frac{a}{2\pi\sigma_2^2} \exp\left(-\frac{x^2+y^2}{2\sigma_2^2}\right)\right)^2 dx\,dy$. The neighbourhood size is then $\mathcal{N} = 1/P_{id} = 4\pi\, N \,/\, \left( \frac{(1-a)^2}{\sigma_1^2} + \frac{a^2}{\sigma_2^2} + \frac{4a(1-a)}{\sigma_1^2+\sigma_2^2} \right)$, where $n \to 4\sigma_l^2 N$ gives the number of potential parents in a square of side $2\sigma_l$ (note that $\sigma_l$ cancels out) and $N$ is the local density, in this IBM approximated by the number of individuals within a deme. For a Gaussian dispersal ($a = 0$), this reduces to $\mathcal{N} = 4\,\pi\,\sigma^2 N$; in the haploid model used here, the neighbourhood size is half as large. Note that while there is no simple expression in terms of $\sigma_{\text{mix}}$ and $\kappa$, the neighbourhood size declines with kurtosis $\kappa$ for a given variance. This contrasts with the formula in [43, Box 2] which seems to be erroneous (and does not appear in [62] as stated in [43]). (C) 10% of dispersal with $\sigma_2 = 4\sigma$ ($\kappa \approx 12.7$, $\sigma^2_{\text{mix}} = 1.25$). For moderate kurtosis and mainly local dispersal, the expansion threshold with adjusted $\mathcal{N}$ and $B$ (dotted green line) appears to work reasonably well. Thus, if we measure $B$ from the experienced fitness cost of dispersal, it should give the appropriate critical rate of temporal change $k^*$. (D) 30% of dispersal with $\sigma_2 = 4\sigma$ ($\kappa \approx 7.7$, $\sigma^2_{\text{mix}} = 2.75$): with an increasing proportion of the wider dispersal kernel, the kurtosis $\kappa$ declines again: yet, in the IBM, the collapse occurs for a slower effective rate of temporal change $k^*$ than expected, notably when the effective gradient $B$ is steep. This is presumably due to excess stochasticity when the simulated reproduction is strictly local, the dispersal is kernel is rather wide and the population shifts fast across steep gradients.

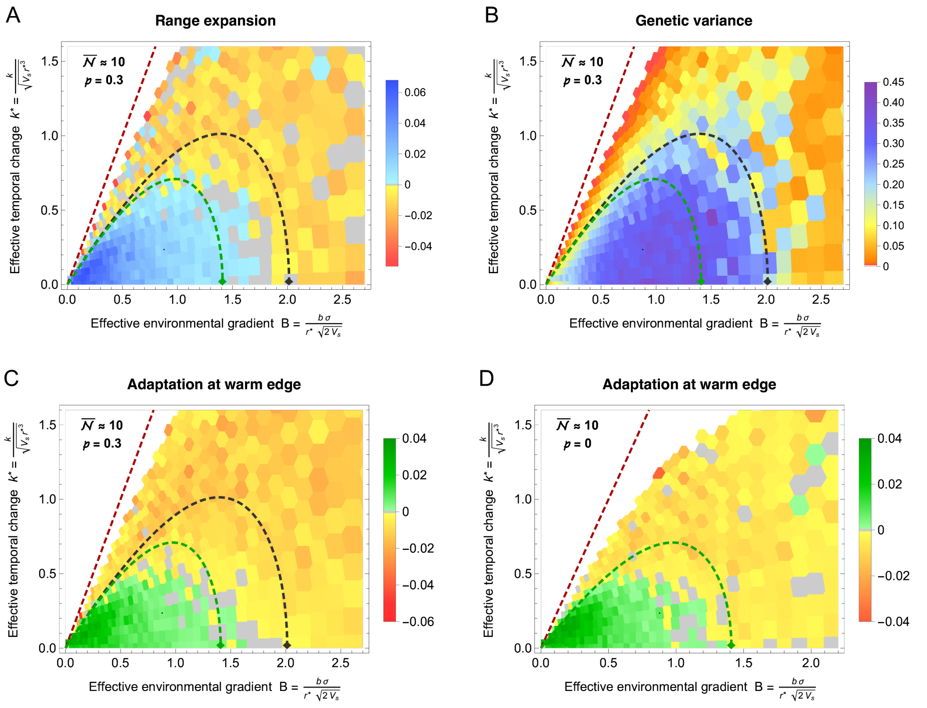


**Fig. S8 Adaptive phenotypic plasticity facilitates adaptation to spatial but not temporal change when genetic variance is maintained by migration-mutation-selection balance**. (A) Species' range continues to expand under steeper effective environmental gradients $B$ in the presence of adaptive phenotypic plasticity $\mathfrak{p}$, which renders the expressed trait $z$ closer to the optimum $\theta$ than expected based on its genetic basis alone, $(z-\theta) \to (1-\mathfrak{p})(z-\theta)$. The effect of plasticity is depicted by the dashed black curve, while the reference without plasticity (as in Fig. 1) is shown in green. (B) Genetic variance is maintained for higher $B$ since the impact of the environmental gradient on fitness, and the associated increase in genetic drift, is reduced. However, the limiting temporal change $k_e^*$ for a given gradient $B$, beyond which adaptation ceases, does not change significantly due to plasticity. This occurs because adaptive plasticity $\mathfrak{p}$ weakens effective selection, thereby reducing the extent to which genetic variance, essential for adaptation to temporal change, increases with the spatial gradient. At the deterministic limit, variance will be reduced by a factor of $(1-\mathfrak{p})$. Hence, the effect of phenotypic plasticity cancels. (C) The rate of adaptation at the *warm* edge (i.e., in the direction of temporal change in the optimum) improves only marginally due to plasticity, and only when the spatial gradient $B$ is steep and the rate of temporal change $k^*$ is slow. (D) The rate of adaptation in the absence of plasticity is similar but shows noticeably higher noise due to greater demographic stochasticity. Parameters are as in Fig. 1, except for plasticity, which is set to $\mathfrak{p} = 0.3$ in (A–C). Median neighbourhood size $\overline{\mathcal{N}} = 2\pi\sigma^2 N \approx 10$, ranging from 8 to 17, and is higher for lower $B$ and $k^*$, where the overall demographic load is lower. Median total population size is approximately $4 \cdot 10^3$ for all surviving populations, ranging from 640 to $3.2 \cdot 10^5$ for expanding populations at $T = 500$ generations.

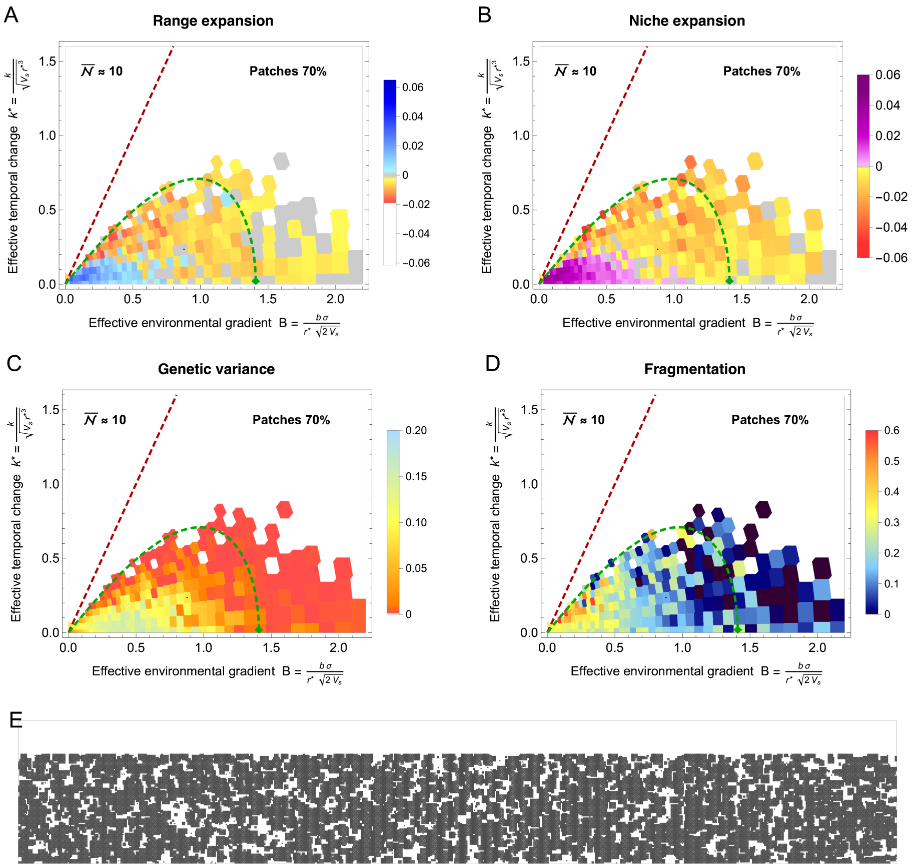


**Fig. S9 Habitat fragmentation hinders adaptation as well as shifts of the species' range.** This figure illustrates the effect of habitat fragmentation, implemented such that the environmental gradient remains linear but the available habitat is patchy (see (E)). (A) With habitat fragmentation, the species' range contracts faster, particularly under rapid temporal change. This is expected because patchiness restricts the ability of the species to track the old optimum through space. However, habitat fragmentation has a negative effect even in the absence of temporal change when the environmental gradient B is steep. (B) Habitat fragmentation substantially restricts adaptation. In particular, in the presence of temporal change, the species' niche contracts faster than in the absence of habitat fragmentation. (C) The lack of adaptation is associated with depleted local genetic variance. (D) Habitat patchiness induces excess range fragmentation, particularly under fast temporal change. Note that for steep gradients, the fragmentation appears to drop again, yet this is not a sign of recovery but rather of the last fragment(s) persisting: with fragmentation of zero, only one fragment is left. This is also clear from the depleted local genetic variance, which implies that there cannot be continuous adaptation along the gradient. (E) An example of 30% habitat fragmentation, generated independently for each run. The mosaic structure arises by adding square habitat patches of side length 8 and a local carrying capacity K until 70% of the space is filled. The median neighbourhood size $\mathcal{N} = 2\pi\sigma^2 N \approx 11$ ranges from 7 to 17, and is higher for lower $B$ and $k^*$, where the overall demographic load is lower. The median total population size is 1124 for all surviving populations – three times smaller than for a continuous habitat; the difference arises because continuous adaptation fails for weaker effective spatial gradients $B$ and slower effective temporal change $k^*$, and these populations also collapse faster in a patchy environment. The median census size is about halved at $T = 500$ generations: 573 *vs.* 1036 for continuous habitat. Expanding populations have a median of $10^5$, ranging from 210 to $2.2 \cdot 10^5$.

**Supplementary Movie S1 Temporal dynamics of species' range fragmentation.**
Top: At time zero, the trait mean (blue line) matches the optimum (grey dashed line). As the optimum begins to shift at the effective rate $k^* = 0.285$, the lag between the trait mean and the optimum increases. This leads to an increase in lag load (the fitness cost of mean maladaptation), which in turn increases genetic drift. Increasing genetic drift erodes clines in allele frequencies and hence genetic variance, which leads to a reduced rate of adaptation, and thus a further increase in the lag load – creating a feedback loop that leads to abrupt range and niche fragmentation. The disjunct subpopulations now track their optima through space (provided the old habitat remains available). Blue dots show individual trait values. Bottom: In the absence of temporal change ($T = 0$, after burn-in with $k^* = 0$), the population is continuous, and its genetic variance is on average close to the expectation for spatially continuous adaptation, $V_G \approx 0.21$. As the lag load and genetic drift increase, variance is depleted, and the species' range fragments abruptly; most of the fragmentation occurs by generation 30. Parameters are the same as in Fig. 4. The first 100 generations are shown; for the purpose of the video, the simulation was rerun, saving the population every generation.
